# Mapping of a N-terminal α-helix domain required for human PINK1 stabilisation, Serine228 autophosphorylation and activation in cells

**DOI:** 10.1101/2021.09.06.459138

**Authors:** Poonam Kakade, Hina Ojha, Olawale G. Raimi, Andrew Shaw, Andrew D. Waddell, James R. Ault, Sophie Burel, Kathrin Brockmann, Atul Kumar, Mohd Syed Ahangar, Ewelina M. Krysztofinska, Thomas Macartney, Richard Bayliss, Julia C. Fitzgerald, Miratul M. K. Muqit

## Abstract

Human autosomal recessive mutations in the *PINK1* gene are causal for Parkinson’s disease (PD). PINK1 encodes a mitochondrial localised protein kinase that is a master-regulator of mitochondrial quality control pathways. Structural studies to date have elaborated the mechanism of how mutations located within the kinase domain disrupt PINK1 function, however, the molecular mechanism of PINK1 mutations located upstream and downstream of the kinase domain are unknown. We have employed mutagenesis studies of human PINK1 in cells to define the minimal region of PINK1, required for optimal ubiquitin phosphorylation, beginning at residue Ile111. Bioinformatic analysis of the region spanning Ile111 to the kinase domain and inspection of the AlphaFold human PINK1 structure model predicts a conserved N-terminal α-helical domain extension (NTE domain) within this region corroborated by hydrogen/deuterium exchange mass spectrometry (HDX-MS) of recombinant insect PINK1 protein. The AlphaFold structure also predicts the NTE domain forms an intramolecular interaction with the C-terminal extension (CTE). Cell-based analysis of human PINK1 reveals that PD-associated mutations (e.g. Q126P), located within the NTE:CTE interface, markedly inhibit stabilization of PINK1; autophosphorylation at Serine228 (Ser228); and Ubiquitin Serine65 (Ser65) phosphorylation. Furthermore, we provide evidence that NTE domain mutants do not affect intrinsic catalytic kinase activity but do disrupt PINK1 stabilisation at the mitochondrial Translocase of outer membrane (TOM) complex. The clinical relevance of our findings is supported by the demonstration of defective stabilization and activation of endogenous PINK1 in human fibroblasts of a patient with early-onset PD due to homozygous PINK1 Q126P mutations. Overall, we define a functional role of the NTE:CTE interface towards PINK1 stabilisation and activation and show that loss of NTE:CTE interactions is a major mechanism of PINK1-associated mutations linked to PD.

## Introduction

Autosomal recessive mutations in PTEN-induced kinase 1 (PINK1) are associated with early-onset Parkinson’s disease (PD) [1]. Human PINK1 (hPINK1) encodes a 581 amino acid Ser/Thr protein kinase containing a N-terminal mitochondrial targeting sequence (MTS) (residues 1-34); catalytic kinase domain containing three major loop insertions (residues 156-510) and a C-terminal extension (CTE) (residues 511-581) [2]. Under basal cellular conditions, PINK1 is imported into mitochondria via its MTS whereupon it undergoes consecutive cleavage by mitochondrial processing protease (MPP) and the rhomboid protease PARL. The N-terminally cleaved PINK1 fragment (residues Phe104-end) subsequently undergoes 20S proteasomal degradation via the N-end rule pathway [3].

Upon mitochondrial depolarisation, that can be induced by mitochondrial uncouplers (e.g. Antimycin A/Oligomycin or carbonyl cyanide m-chlorophenyl hydrazone (CCCP)), full-length human PINK1 protein is stabilised at the mitochondria where it becomes autophosphorylated and active [4]. Active PINK1 phosphorylates ubiquitin at Serine (Ser65) that recruits the Parkinson’s linked ubiquitin ligase, Parkin, to the mitochondrial surface whereupon it phosphorylates Parkin at an equivalent Ser65 residue within its N-terminal Ubiquitin-like (Ubl) domain leading to activation of Parkin E3 ligase activity and ubiquitin-dependent elimination of damaged mitochondria by autophagy (mitophagy) [5-7]. Active PINK1 also indirectly induces the phosphorylation of a subset of Rab GTPases including Rab 8A at a highly conserved Serine residue (Ser111) that lies within the RabSF3 motif [8].

Structural and biochemical analysis of catalytic domain-containing fragments of insect orthologues of PINK1 namely, *Tribolium castaneum* (Tc) [PDB 5OAT] and *Pediculus humanus corporis* (Phc) (in complex with Ubiquitin via a nanobody) [PDB 6EQI], have shed light on intrinsic kinase domain-mediated mechanisms of PINK1 activation [9, 10]. These studies identified a critical role of autophosphorylation of a Serine residue (TcPINK1, Ser205; PhcPINK1 Ser202) within the N-lobe for activation and ubiquitin substrate recognition that is equivalent to Ser228 of hPINK1 [9, 10]. Structural analysis also revealed a previously unidentified role for the third loop insertion (INS3) towards substrate recognition and catalytic activity [9, 10]. Mutagenesis studies of human PINK1 in cells has confirmed conservation of these mechanisms however, unambiguous detection of PINK1 autophosphorylation at Ser228 (equivalent to the insect pSer205/202) is outstanding.

The structures have provided atomic insights into the pathogenic mechanism of key human PINK1 disease-associated mutations that lie within the kinase domain. However, a number of disease mutations lie outside the kinase domain within the upstream N-terminal region of PINK1 and current published structures provide no insights into its function [9, 10]. Upon mitochondrial depolarisation, full-length human PINK1 accumulates on the outer mitochondrial membrane (OMM) in association with the Translocase of outer membrane (TOM) complex, however, the molecular basis of how human PINK1 is stabilised and activated at the TOM complex remains to be fully elucidated [11, 12].

Herein we have mapped the minimal region of human PINK1, required for optimal ubiquitin phosphorylation, beginning at residue Ile111. Bioinformatic analysis of the region spanning Ile111 and the kinase domain and inspection of a predicted structural model of human PINK1 by AlphaFold [13] reveals a N-terminal α-helical domain extension (NTE domain) which forms intramolecular contacts with the CTE. Mutagenesis of key residues at the NTE-CTE interface indicate a critical role of this interface towards human PINK1 stabilisation at the TOM complex; Ser228 autophosphorylation and PINK1 activation in cells.

## Materials and Methods

### Reagents

[γ-^32^P] ATP was from PerkinElmer, MLi-2 was obtained from Merck [14]. LRRK2 (G2019S) Recombinant tetra-ubiquitin was generated by Dr Axel Knebel (MRC PPU). All mutagenesis was carried out using the QuikChange site-directed mutagenesis method (Stratagene) with KOD polymerase (Novagen). cDNA constructs for mammalian tissue culture (Supplementary Table 1) were amplified in Escherichia coli DH5α and purified using a NucleoBond Xtra Midi kit (#740420.50; Macherey-Nagel). All DNA constructs were verified by DNA sequencing, which was performed by The Sequencing Service, School of Life Sciences, University of Dundee, using DYEnamic ET terminator chemistry (Amersham Biosciences) on Applied Biosystems automated DNA sequencers. DNA for bacterial protein expression was transformed into *E. coli* BL21 DE3 RIL (codon plus) cells (Stratagene). All cDNA plasmids (Supplementary Table 1), CRISPR gRNAs, antibodies and recombinant proteins generated for this study are available to request through our reagents website https://mrcppureagents.dundee.ac.uk/.

### Cell Culture and Transfection

SKOV3 and HeLa WT & PINK1 knock-out cells were routinely cultured in standard DMEM (Dulbecco’s modified Eagle’s medium) supplemented with 10% FBS (foetal bovine serum), 2 mM L-Glutamine, 100 U/ml Penicillin & 100 mg/ml Streptomycin (1X Pen/Strep), and 1X non-essential amino acids (Life Technologies). Flp-In™ T-REx™ HEK293 cells & HeLa cells were supplemented with 15 μg/ml of Blasticidin and 100 μg/ml of Hygromycin and were induced to express protein by addition of 0.1 μg/ml of Doxycycline to the medium for 24 h. All cells cultured at 37°C, 5% CO2 in a humidified incubator and routinely tested for Mycoplasma.

For transient expression, SKOV3 cells were transfected at 90% confluency using Lipofectamine 3000 (3.2 μL Lipofectamine and 4 μL P3000 reagent used per 10 cm plate). Cells were then cultured for a further 72 hrs including treatment of Oligomycin and Antimycin A for 9 h. HeLa cells were transiently transfected with 4-5 μg of DNA dissolved in 2 mL serum-free DMEM, mixed with 60 μL 1mg/mL PEI, vortexed gently for 10 sec and incubated for 20-40 min at room-temperature before being added dropwise to cells at around 50-60% confluency. To uncouple mitochondria, cells were treated with or without 10 μM CCCP (Sigma) dissolved in DMSO for 6 h or 10 μM Antimycin A / 1 μM Oligomycin for 3 h. Cells were harvested in ice-cold lysis buffer as explained below.

### Generation of CRISPR-Cas9 PINK1 knockout HEK293 cells and HeLa cells

CRISPR was performed using a paired Nickase approach to minimize off-target effects. Exon 2-specific guide pairs with low combined off-targeting scores were identified using the Sanger Institute CRISPR webtool (http://www.sanger.ac.uk/hgt/wge/find_crisprs). Complementary oligos for the optimal guide pair A (G1 5’-gCTTGCAGGGCTTTCGGCTGG and G2 5’-gCGTCTCGTGTCCAACGGGTC) were designed and annealed according to the Zhang method with BbsI compatible overhangs facilitating cloning into the target vectors; the sense guide G1 was cloned into the puromycin selectable plasmid pBABEDPU6 (DU48788 https://mrcppureagents.dundee.ac.uk/) and antisense guide G2 cloned into the spCas9 D10A-expressing vector pX335 (Addgene Plasmid #42335) yielding clones DU52528 and DU52530, respectively. CRISPR was performed by co-transfection of Flp-In™ T-REx™ HEK293 cells/ Flp-In™ T-REx™ HeLa cells (60% confluency, 10cm dish) with 3.75 μg of each plasmid in 27 μl Lipofectamine LTX according to manufacturer’s instructions. Following CRISPR, media was removed and transfected cells were selected by twice incubation for 48 hrs in complete media supplemented with 3 μg/ml puromycin. Following the second incubation, cells were returned to fresh media lacking puromycin for recovery of selected cells. Single cells were isolated by flow-assisted cell-sorting into individual wells of a 96-well plate containing DMEM supplemented with 10% FBS, 2mM L-Glutamine, 100 U/ml Penicillin, 100 μg/ml Streptomycin and 100 μg/ml Normocin (InvivoGen). After reaching ∼80% confluency, individual clones were transferred into 6-well plates and PINK1 expression determined by immunoblot analysis following CCCP treatment. Knockouts were confirmed by sequencing.

Validated PINK1-knockout Flp-In™ T-REx™ HEK293clone “F1” and Flp-In™ T-REx™ HeLa clone “3C9” were utilised as the parental lines for the generation of Doxycycline-inducible, PINK1-3FLAG stably-expressing cells. Cell lines were generated according to manufacturer’s instructions by selection with hygromycin. Expression was confirmed by immunoblot following treatment with 20 ng/ml Doxycycline.

### Generation of Flp-In™ T-REx™ PINK1-rexpression stable cell lines

To ensure low-level uniform expression of recombinant proteins, manufacturer’s instructions (Invitrogen) were followed to generate stable cell lines that re-express C-terminal 3FLAG-tagged forms of PINK1 proteins (cDNA subcloned into pcDNA5-FRT/TO plasmid) in a Doxycycline inducible manner. Flp-In T-REx-293 / Flp-In T-rex-Hela CRISPR knock-out to PINK1 null cells were generated in lab. The PINK1 null host cells containing integrated FRT recombination site sequences and Tet repressor, were co-transfected with 4.5/9 μg of pOG44 plasmid (which constitutively expresses the Flp recombinase), and 0.5/1 μg of pcDNA5-FRT/TO vector containing a Hygromycin resistance gene for selection of the gene of interest with FLAG tag under the control of a Doxycycline-regulated promoter. Cells were selected for Hygromycin and Blasticidin resistance three days after transfection by adding fresh medium supplemented with 15 μg/ml of Blasticidin and 100 μg/ml of Hygromycin. Expression of the recombinant protein was induced by addition of 0.1 μg/ml of Doxycycline for 24 hours.

### Primary human skin fibroblasts

Primary skin fibroblasts at low passage numbers (2–5) were contributed by the Hertie Institute for Clinical Brain Research Biobank, Tübingen, Germany. They were obtained from skin biopsies from a patient with PD with a PINK1 p.Q126P homozygous mutation (p.Q126P hom; c.388-7A>G hom; c.960-5G>A hom; c.*37A>T hom) [15] and age-matched healthy individuals following routine clinical procedures, underwritten informed consent (Hertie Institute for Clinical Brain Research Biobank) and approval by a local ethics committee (University of Tübingen). Patients were screened for GBA, LRRK2, PARK2, DJ-1 and PINK1 mutations by RFLP, MPLPA and bidirectional Sanger sequencing of the entire coding sequence of PINK1 using an ABI PRISM 3100-Avant Genetic Analyzer (Applied Biosystems), as described previously [15]. Fibroblasts were cultured in Dulbecco’s modified Eagle’s medium supplemented with glucose (4.5 g/l), L-glutamine (2 mM), HEPES(10 mM), foetal bovine serum (10%) and penicillin (50 U/ml)/streptomycin (50 lg/ml) plus 1% (v/v) non-essential amino acid and grown at 37°C in a 5% CO2 atmosphere.

### Whole cell lysate preparation

Cells were lysed in an ice-cold lysis buffer containing 50 mM Tris–HCl, pH 7.45, 1% (by vol) Triton X-100, Sucrose 250 mM, 1mM EDTA, 1mM EGTA, 1 mM sodium orthovanadate, 50 mM NaF, 10 mM 2-glycerophosphate, 5 mM sodium pyrophosphate, and complete EDTA-free protease inhibitor cocktail (Roche) with freshly added 1x phosphatase inhibitor cocktail (Sigma-Aldrich). Lysates were clarified by centrifugation at 16,600 g at 4°C for 20 min and supernatants were quantified by BCA assay.

### Isolation of mitochondrial enriched fraction

Cells were collected in ice-cold mitochondria fractionation buffer containing 20mM HEPES pH 7.5, 3 mM EDTA, 5 mM Sodium β-glycerophosphate, 50 mM Sodium fluoride, 5 mM Sodium pyrophosphate, 250 mM sucrose, 1mM Sodium orthovanadate, 1x protease inhibitor cocktail (Roche), and 200mM chloroacetamide. Cell pellets were disrupted by 25 passes in a handheld homogeniser, or 25-gauge needle, and lysates clarified by centrifugation at 800 g, 10 min at 4 °C. Supernatant was isolated and centrifuged at 16,000 g for 10 min at 4 °C. The resultant supernatant was retained as cytoplasmic fraction and the pellet was resuspended in mitochondria fractionation buffer with 1% Triton X-100 and retained as mitochondrial-enriched fraction. Samples were quantified by Bradford and normalised appropriately.

### Blue Native PAGE (BN-PAGE)

The samples for BN-PAGE analysis were prepared using a Native PAGE Sample Prep Kit (Invitrogen). For BN-PAGE, mitochondria-enriched fractions were gently pipetted up and down 10 times in 1× NativeTM PAGE buffer with 1% digitonin followed by an incubation for 30 min at 4 °C. The samples were centrifuged at 20,000 g for 30 min at 4 °C. Samples were quantified by BCA assay and supplemented with 0.002% G-250 (Invitrogen). BN-PAGE was performed by NativeTM PAGE Running Buffers (Invitrogen). The gels were washed in denaturation buffer (10 mM Tris-HCl pH 6.8, 1% SDS, and 50mM DTT) for 15 min at 60 °C after electrophoresis and then transferred on to PVDF membranes for IB analysis. PVDF membranes were incubated in destaining buffer (40% methanol, 10% acetic acid) and subjected to immunoblotting.

### PINK1 immunoprecipitation

For immunoprecipitation of endogenous PINK1, 500 µg of whole-cell was incubated overnight with PINK1 antibody (S085D, MRC PPU reagents and Services) coupled to Protein A/G beads as previously reported [16]. The immunoprecipitants were washed three times with lysis buffer containing 150 mM NaCl and eluted by resuspending in 10 µl of 2× LDS sample buffer and incubating at 37°C for 15 min under constant shaking (2000 rpm) followed by the addition of 2.5% (by vol) 2-mercaptoethanol.

### Ubiquitin Pulldown

For ubiquitylated protein capture, Halo-tagged ubiquitin-binding domains (UBDs) of TUBE (tandem-repeated ubiquitin-binding entities) was incubated with HaloLink resin (200 μL, Promega) in binding buffer (50 mm Tris⍰HCl, pH 7.5, 150 mm NaCl, 0.05 % NP-40) overnight at 4 °C. Membrane-enriched fraction (400 μg) was used for pulldown with HALO-UBDs. Halo Tube beads (20 μL) were added to whole cell lysates for enrichment and incubated O/N at 4 °C. The beads were washed three times with lysis buffer containing 0.25 mM NaCl and eluted by resuspension in 1×LDS sample buffer (20 μL) with 1 mM dithiothreitol (DTT) or 2.5% 2-mercaptoethanol. The method for Ubiquitin capture using has previously been reported for Halo-tagged ubiquitin-binding domains (UBDs) of TUBE (tandem-repeated ubiquitin-binding entities) [17].

### Immunoblotting

Samples were subjected to SDS-PAGE and transferred onto nitrocellulose membranes. Membranes were blocked with 5 % BSA in TBS-T and incubated at 4 °C overnight with the indicated antibodies, diluted in 5 % BSA. Membranes for kinase assay screen were incubated with HRP-conjugated secondary antibodies (1:10,000) in 5 % milk for 1 h at room temperature. Membranes were exposed with ECL substrate. Additional kinase assay immunoblotting membranes were incubated with secondary antibodies, conjugated with LICOR IRDye in TBS-T (1:10,000) and imaged using the LICOR Odyssey software.

### Antibodies

The following primary antibodies were used: Parkin phospho-Ser65 rabbit monoclonal antibody was raised by Epitomics/Abcam in collaboration with the Michael J Fox Foundation for Research (Please contact for questions). Ubiquitin phospho-Ser65 (Cell Signalling Technology (CST)), OPA1 (CST), PINK1 (Novus), Parkin (Santa Cruz), Rab8A (Abcam), Rab8A phospho-Ser111 (Abcam), GAPDH (Santa Cruz), and Vinculin (CST). The polyclonal phospho-Ser228 PINK1 was generated by the Michael J. Fox Foundation’s research tools program in partnership with Abcam (Development of a monoclonal antibody is underway. Please contact for questions). The following antibody was produced by the MRCPPU Reagents and Services at the University of Dundee in sheep: anti-PINK1 (S085D).

### Expression of recombinant Pediculus PINK1 proteins

Wild-type (WT) and mutant *Pediculus humanus corporis* PINK1 (Phc) (residues 108-end) were expressed as a His-SUMO tagged protein with SENP1 cleavage site in BL21(DE3)pLysS. Cells were grown in Terrific broth medium containing 50 µg ampicillin to an OD of 0.8 and expression was induced at 16 °C with 250 µM IPTG and allowed to grow for a further 16 h. Cells were harvested by centrifugation at 4000 rpm for 30 min, cell pellet was resuspended in lysis buffer; 25 mM Tris pH 8.5, 300 mM NaCl, 0.5 mM TCEP, 5% glycerol (containing lysozyme, DNAase and protease inhibitor cocktail and 10 mM imidazole). Cells were lysed by sonication and cell lysate centrifuged using ultra centrifuge at 20000xg to collect cell debris. The supernatant was incubated with nickel beads at 4 °C for 2 h on a rotatory platform. After 2 h, beads were collected and washed using lysis buffer containing 10 mM imidazole. Protein was eluted using lysis buffer containing 200 mM imidazole. His-SUMO was cleaved off overnight by dialysing into dialysis buffer (25 mM Tris, 300 mM NaCl pH 8.5, 0.5 mM TCEP, 5% glycerol) using His-SENP1 at 4 °C. Cleaved protein was passed through fresh nickel beads to remove His-SUMO and His-SENP1. Eluted protein was collected, concentrated and purified further by gel filtration using gel filtration buffer(25mM Tris, pH8.5, 150mM NaCl, 5% glycerol and 0.5mM TCEP). The elution profile of the protein was monitored using a calibrated Superdex 75 column. Pure fractions from SDS-PAGE analysis were collected, concentrated, quantified and flash frozen until needed.

### Pediculus PINK1 *in vitro* kinase assay

*In vitro* activity assays were set up in a final volume of 40 μl containing 2 μM substrate (K63-linked Tetra-Ub) in 50 mM Tris-HCl pH 7.5, 0.1 mM EGTA, 10 mM MgCl2, and 0.1 mM [γ-32P]ATP (approximately 500 cpm/pmol). 250 nM of Phc-PINK1 (WT) and mutants as kinase was used. Assays were incubated at 30 °C for 10 min with shaking at 1050 rpm, and terminated by addition of 13 μl 4x LDS sample buffer containing reducing agent. Samples were boiled at 70 °C for 10 min and reactions resolved by SDS-PAGE. For autoradiography, gels were stained for protein detection for 1 hr in Coomassie InstantBlue, then destained by washes in warm water. Wet gels were scanned in an Epson scanner, then dried completely using a gel dryer (Bio-Rad). Incorporation of [^γ32^P]ATP into substrates was analysed by exposure to Amersham Hyper-Film at -80 °C. For assay quantification, individual SDS-PAGE bands were excised from dried gels and incorporation determined by Cerenkov counting. Kinase activity is determined by quantification of the number of phosphates transferred by a kinase, per minute of the reaction, per kinase molecules in the reaction. Number of phosphates transferred is determined by relating the scintillation counts of a substrate band to the counts of a known amount of phosphate i.e. the 1mM [^y-32^P]ATP stock.

### Intact mitochondrial kinase assay (MitoKA)

Intact mitochondrial assay was performed as previously described [18]. Intact mitochondria-enriched fractions were isolated in CFAB buffer (CFAB: 20 mM HEPES KOH (pH 7.5), 220 mM sorbitol, 10 mM potassium acetate, 70 mM sucrose protease inhibitor cocktail minus EDTA (Roche)) were incubated with 2 μM substrate (tetra ubiquitin (K63 linked)) 2 mM ATP, 2 mM DTT, 5 mM MgCl2 and 1% glycerol (50 μl final) for indicated time points. Reactions stopped by addition of 16.6 μl 4x LDS 4% 2-beta-mercaptoethanol and samples were vortexed and heated at 95°C for 5 min. Samples were diluted 1:10 in 1x LDS for substrate immunoblotting and remainder used for immunoblotting controls and Coomassie staining.

### Hydrogen/deuterium exchange mass spectrometry (HDX)

The HDX experiment was performed as described by Cornwell *et al*., (2018) with modifications [19]. Briefly, 5 µL of 5 µM of wild type PhcPINK1 (residues 108-575) was equilibrated in 50 mM potassium phosphate, 300 mM sodium chloride, 5 mM TCEP, pH7.2. Sample was mixed with 95 µl deuterated buffer (10 mM potassium phosphate, 300 mM sodium chloride, 5 mM TCEP pD 6.8), and was then incubated at 4 °C for the specified times (0.5 min or 2 min). At the end of the labelling time 100 µL of the reaction mixture was added to 100 µL of the quench buffer at 0 °C (10 mM potassium phosphate, 5 mM TCEP, 0.1% DDM. The quench buffer pH was adjusted so the final mixture was at pH 2.5). 50 µl of the quenched sample was passed through an immobilised pepsin column (Affipro, Prague, Czech Republic) at 150 µl/min at 20 °C and the resulting peptides were trapped by a VanGuard Acquity UPLC BEH C18 pre-column for 3 min. The peptides were then transferred to a C18 column (75 µm × 150 mm, Waters Ltd., Wilmslow, Manchester, UK) and separated by gradient elution of 5-40% MeCN in water and 0.1% formic acid over 7 min at 40 µl/min. Trapping and gradient elution of peptides was performed at 0M°C.

The HDX system was interfaced to a Synapt G2Si mass spectrometer (Waters Ltd., Wilmslow, Manchester, UK). HDMS^E^ and dynamic range extension modes (Data Independent Analysis (DIA) coupled with IMS separation) were used to separate peptides prior to CID fragmentation in the transfer cell [20]. Data was analysed using PLGS(V3.0.2) and DynamX(v3.0.0) software which were supplied with the mass spectrometer as previously described [19].

## Results

### Mitochondrial import and association via the TOM complex is required for PINK1 activation

It has previously been reported that stable expression of PINK1 at the outer mitochondrial membrane (OMM) using a fusion construct of N-terminally truncated PINK1 with the OMM anchor sequence of OPA3 (OPA3 (1-30)-PINK1 (111-end) is sufficient to activate human PINK1 independent of mitochondrial depolarisation as determined using the Parkin-mitochondria recruitment assay [21]. We initially explored whether PINK1 stabilisation at the OMM or association with the TOM complex is sufficient for its activation as measured by monitoring phosphorylation of its substrates Parkin and ubiquitin. We first generated Flp-In™ T-REx™ HEK293 PINK1-knockout cells by exon 2-targeted CRISPR-Cas9 (Supplementary Figures 1A-B), then stably re-introduced full-length PINK1-3FLAG wild-type (WT); kinase-inactive mutant (KI; D384A); OPA3 (1-30)-PINK1111-581 (OPA3-111PINK1); or PINK1 fused to a conventional N-terminal MTS that is imported to the matrix via the TOM, namely HSP60 (1-27)-PINK1111-581 (HSP60-111PINK1) (Figure 1A). Cells were treated with DMSO or 10 μM CCCP for 3 h, and following cell lysis, mitochondrial extracts were analysed by immunoblotting with anti-FLAG antibody (Figure 1B). This confirmed that HSP60-111PINK1 underwent proteolytic cleavage under basal conditions indicating successful import to the mitochondrial matrix and this was reduced upon mitochondrial depolarisation (Figure 1B). In contrast, OPA3-111PINK1 was stably expressed at the mitochondria with no evidence of proteolysis under basal conditions and no change upon mitochondrial depolarisation, suggesting that it is stabilized at the OMM and not imported (Figure 1B). Cells were next transfected with untagged Parkin and treated with DMSO or AO for 9 h. Cells were lysed and whole cell extracts analysed by immunoblotting with anti-phospho-Ser65-ubiquitin, and anti-phospho-Ser65-Parkin antibodies (Figure 1C). Under basal conditions we observed robust stable levels of OPA3-111PINK1 however, this was not active under our assay conditions and furthermore we did not observe any activation of OPA3-111PINK1 following mitochondrial depolarisation (Figure 1B). In contrast we observed that HSP60-111PINK1 was robustly activated upon mitochondrial depolarisation with induction of Ser65-phosphorylated-ubiquitin (phospho-ubiquitin) and Ser65-phosphorylated-Parkin (phospho-Parkin) similar to wild-type PINK1 (Figure 1C).

**Figure 1.**
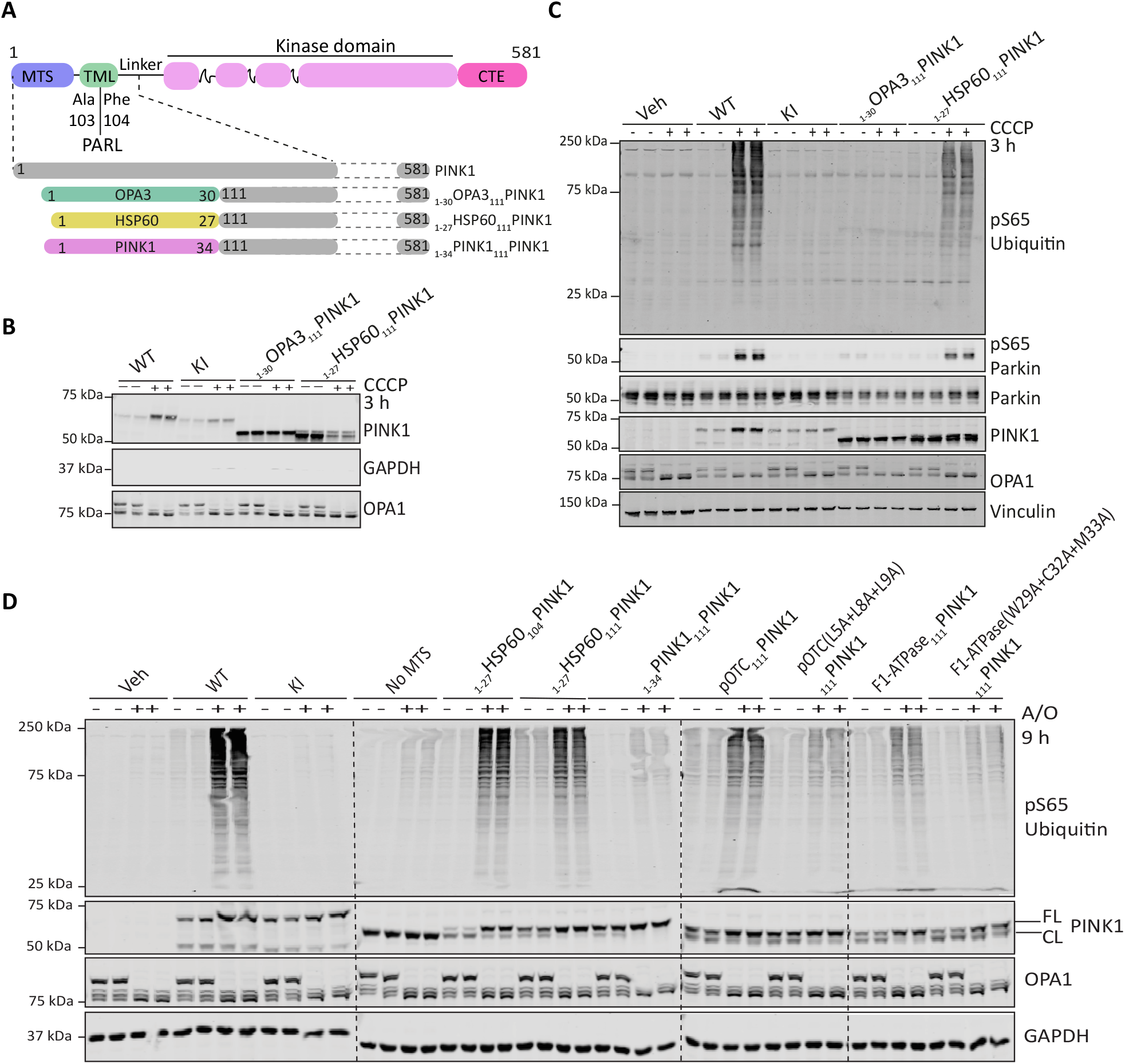
Mitochondrial targeting sequences that mediate TOM complex association are required for PINK1 activation. **A:** Schematic depiction of constructs for PINK1 mutants. For PINK1 mutants targeted to different compartments of the mitochondria, residues 1-110, which includes the mitochondrial targeting sequence (MTS) and transmembrane-like domain (TML) and PARL cleavage site of PINK1, were removed and replaced with the predicted MTS of proteins known to localise to the OMM (OPA3 1- 30) and matrix (HSP60 1-27). For PINK1 mutant with predicted MTS of PINK1, residues 1-110 were replaced with PINK 1 residues 1-34. **B:** Flp-In TRex HEK293 cells PINK1 KO ClF1 stably expressing WT, KI (D384A), OPA3 (1-30)-PINK1(111- 581) or HSP60 (1-27) PINK1(111-581) were treated -/+ 10µM CCCP for 3 hours and subjected to sub cellular fractionation to obtain membrane enriched fraction. Samples were resolved by SDS-PAGE. Proteins were transferred to nitrocellulose membranes probed using the antibodies indicated. n=2. **C:** Flp-In TRex HEK293 cells PINK1 KO stably expressing 3xFLAG alone (Veh), PINK1 wild type (WT), kinase inactive (KI, D384A), OPA3 (1- 30)-PINK1(111-581), and HSP60 (1-27)-PINK1(111-581) were treated with -/+ 10µM CCCP for 3 hours. Whole cell lysates (Parkin and control blots) or membrane enriched fractions (FLAG and pSer^65^ ubiquitin blots) were resolved by SDS PAGE. Proteins were transferred to nitrocellulose membranes probed using the antibodies indicated. **D:** Human SK-OV-3 PINK1 knockout cells were transfected with 3xFLAG alone (Veh), PINK1 wild type (WT), kinase inactive (KI, D384A), HSP60 (1-27)- PINK1(104-581), HSP60 (1-27)-PINK1(111-581), PINK1(1-34)-PINK1(111- 581), pOTC(1-30)-PINK1(111-581) and pF1ATPase(1-30)- PINK1(111-581). Cells were treated with A/O for 9 h prior to lysis and probed using the indicated antibodies.

To confirm these results we utilized human SK-OV-3 ovarian cancer cells that we have identified as a robust system to study endogenous PINK1-Parkin signalling (Wu et al., manuscript under preparation). We performed transient expression of WT PINK1, KI PINK1 and HSP60-111PINK1 in SK-OV-03 PINK1-knockout cells followed by treatment with or without AO for 9 h and observed induction of phospho-ubiquitin in cells expressing HSP60-111PINK1 upon mitochondrial depolarisation consistent with results in Flp-In™ T-REx™ HEK293 cells (Figure 1D). The activation of PINK1 was critically dependent on MTS-directed import since expression of 111PINK1 without a MTS was not activatable (Figure 1D). Furthermore, we did not observe any difference in activation between HSP60-104PINK1 and HSP60-111PINK1. We next tested the generality of our findings using two well studied MTS sequences, pOTC (1-30) and pF1-β ATPase (1-30) which have been shown to mediate mitochondrial import via TOM association [22]. Upon mitochondrial depolarisation, we observed that both pOTC-111PINK1 and pF1-β ATPase-_111_PINK1 led to activation and induction of phospho-ubiquitin. In contrast, mutant sequences of pOTC (L5A/L8A/L9A) and pF1-β ATPase (W29A/C32A/M33A) that have been shown to impair TOM complex interaction [22], were associated with significantly reduced PINK1 activation and phospho-ubiquitin following mitochondrial depolarisation (Figure 1D). We have previously shown that the PINK1 MTS (residues 1-34) is sufficient to localise PINK1 to mitochondria [23]. Interestingly we observed that PINK1[1-34]-111PINK1 was imported and cleaved under basal conditions, however, we did not observe any activation following mitochondrial depolarisation suggesting that additional residues within its N-terminus are required to stabilize it within the TOM complex for activation (Figure 1D). This is consistent with previous studies that have suggested that the sequence determinants for PINK1 import in the native protein are complex, redundant and may include additional MTS sequences located between residues 35-90 [24], and in future studies it will be interesting to define those sites required for activation.

### Mapping the minimum boundary of PINK1 required for activation in cells

To better define the regions of human PINK1 required for optimal activation and phosphorylation of ubiquitin, we initially expressed a series of PINK1 constructs in SK-OV-3 PINK1 KO cells including HSP60-130PINK1, HSP60-125PINK1, HSP60-120PINK1, HSP60-115PINK1, HSP60-111PINK1, and HSP60-104PINK1 (Figure 2A). Immunoblotting analysis with anti-PINK1 antibodies indicated that all PINK1 constructs were expressed at similar levels, however, upon AO treatment, we only observed robust activation of PINK1 in cells expressing either HSP60-104PINK1 or HSP60-111PINK1 that paralleled WT PINK1 and endogenous PINK1 activation (Figure 2B). Since HSP60-115PINK1 was not sufficient for PINK1 activation, we performed fine mapping of the region between residue Ile111 and Gln115 and expressed a series of further constructs including HSP60-114PINK1, HSP60-113PINK1, HSP60-112PINK1, and HSP60-111PINK1. All constructs were expressed at similar levels and underwent cleavage and processing under basal conditions followed by stabilization of the upper PINK1 band upon mitochondrial depolarisation (Figure 2C). However, there was a strict requirement for the N-terminal region of PINK1 to start at residue Ile111 (HSP60-111PINK1) for subsequent activation by mitochondrial depolarisation (Figure 2C). The region of PINK1 between Ile111 and the kinase domain, spanning residues 111-156, is colloquially referred to as the PINK1 “linker’ region and no structural information for this region is currently published. A previous study found that triple mutation of a cluster of negatively charged glutamic acid residues (Glu112Ala/Glu113Ala/Glu117Ala) in this region disrupted accumulation of PINK1 in depolarized mitochondria and its subsequent activation and recruitment of Parkin [Ref/Sekine]. We initially assessed the role of these amino acids in the activation of HSP60-PINK1111-581, by mutating each Glu residue to alanine, however, these did not significantly impact PINK1 activation (data not shown). We next substituted the charge state of each Glu to a Lys residue and expressed these variants in PINK1 KO SK-OV-3 cells. All constructs were expressed at similar levels and underwent cleavage and processing under basal conditions followed by stabilization of the upper band upon mitochondrial depolarisation (Figure 2D). However, the Glu117Lys (E117K) mutation was sufficient to completely prevent PINK1 activation whereas there was no effect of mutation of the other Glu residues including the double Glu112K/Glu113K mutant (Figure 2D).

**Figure 2.**
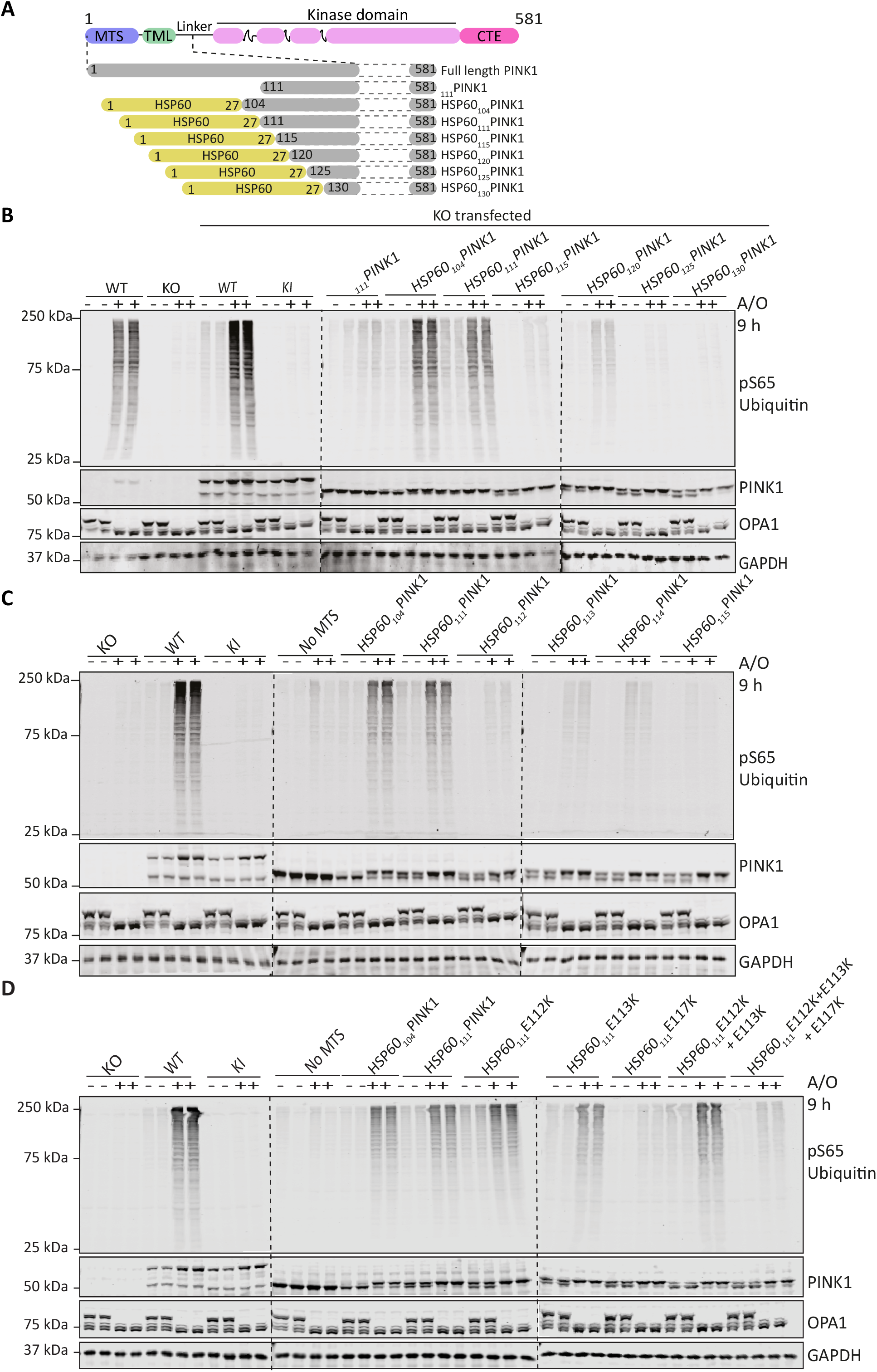
Mapping of N-terminal boundary of minimal region required for PINK1 activation to residue Ile111. **A:** Schematic depiction of different deletion mutants of PINK1 constructed with HSP60-MTS aa 1-27 followed by the region 104-130 to END. **B:** Assay using Human SKOV3 PINK1 WT and PINK1 knock-out (PINK1 KO) cells in combination with PINK1-KO cells were transfected with indicated deletion mutants of PINK1 i.e., HSP60(1-27)-PINK1(104-581) till HSP60(1-27)- PINK1(130-581) along with Wild Type (WT) and Kinase Inactive (KI) were treated with A/O for 9 hrs prior to lysis and immunoblotted with indicated antibodies (anti-pSer65 ubiquitin, anti-PINK1, anti-OPA1 and anti-GAPDH). The membranes were developed using the LI-COR Odyssey CLx Western blot imaging system. **C:** Human SK-OV-3 PINK1 knockout cells were transfected with deletion mutants i.e., HSP60(1-27)-PINK1(111-581) till HSP60(1-27)-PINK1(115-581) cells were treated with A/O for 9 h prior to lysis and indicated antibodies and analysed as described above. **D:** I111 and E117 are crucial for PINK1 dependent phosphorylation of Ubiquitin at Ser65. Human SK-OV-3 PINK1 knockout cells were transfected with E-K and PD mutations of point mutations of the region 111-117 of HSP60(1-27)-PINK1(111-581) till HSP60(1-27)-PINK1(115-581) cells were treated with A/O for 9 h prior to lysis and indicated antibodies and analysed as described above.

### Structural modelling predicts N-terminal α-helix extension (NTE) domain upstream of kinase domain of PINK1 that interacts with CTE

Current insect structures do not include the PINK1 “linker” region spanning residues 111-156 (Figure 3A) [9, 10]. We therefore subjected the linker regions of human PINK1 and PhcPINK1 to secondary structure prediction utilizing SYMPRED [25]. For human PINK1, secondary structure prediction tools consistently return an α-helical structure, reaching a consensus for residues Ile111 - Lys135 with a loop linking the α-helix to the kinase domain (Figure 3B and Supplementary Figures 2A-B). A similar consensus is reached for PhcPINK1 with tools predicting an α-helix from Lys120 - Trp140 followed by a poorly defined region linking to the kinase domain (Figure 3B and Supplementary Figure 2A, C). Furthermore, we analysed the structural model of human PINK1 using AlphaFold [13], and this predicted with high confidence the presence of the N-terminal α-helix extension (NTE) domain extending to Lys135 (Figures 3C-E and Supplementary Figures 2D). These regions are highly conserved within the overall PINK1 sequence (Supplementary Figure 3).

**Figure 3.**
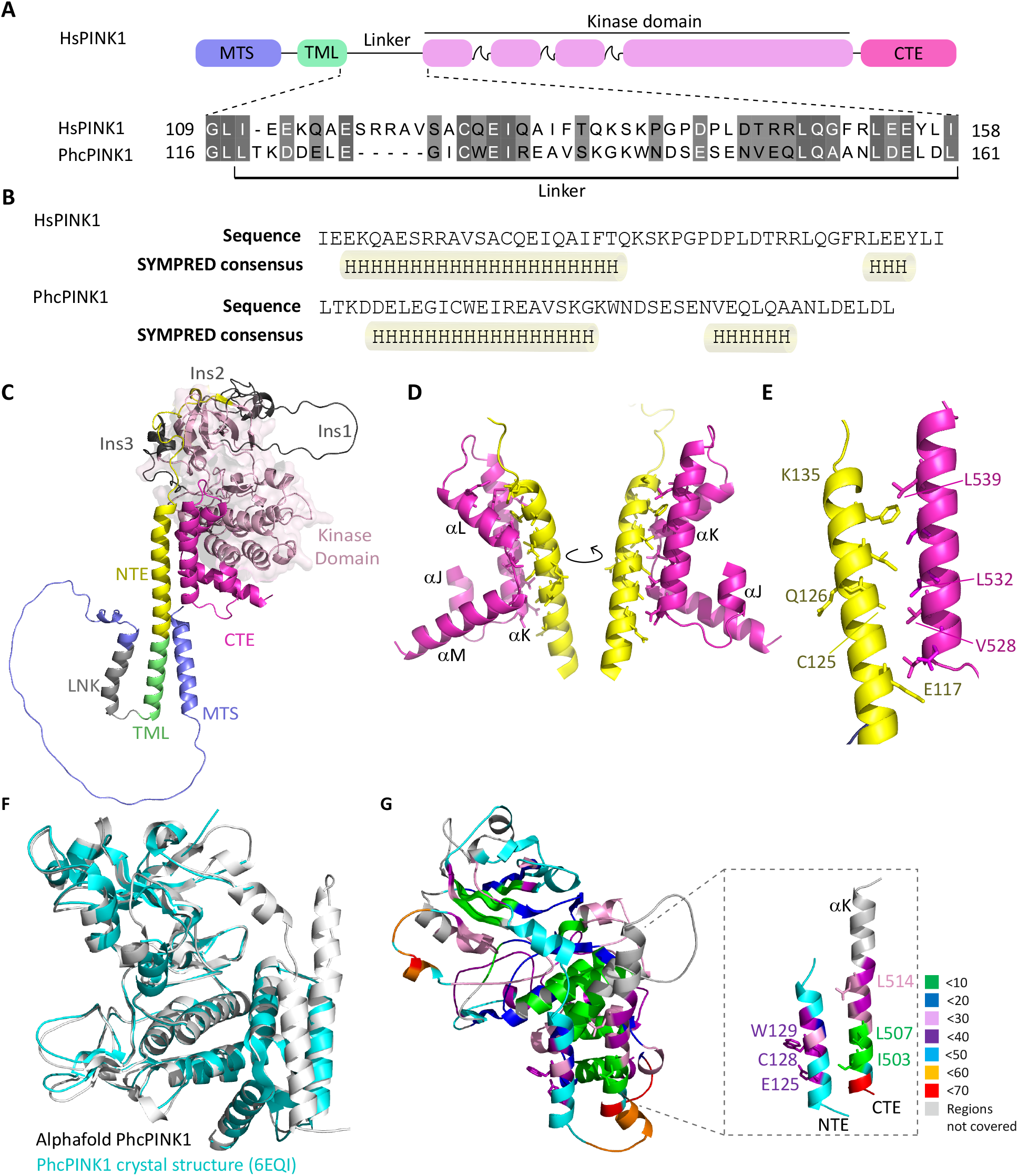
Structural modelling predicts an N-terminal α-helix extension (NTE domain) and its interaction with CTE domain. **A:** Consistent prediction of α-helix in the linker region of PINK1 flanked by transmembrane domain and kinase domain. Schematic representation of human PINK1 (HsPINK1) and Pediculus humanis corporis PINK1 (PhcPINK1) highlighting the linker region flanked by transmembrane domain and kinase domain. The alignment of linker region is performed by MAFT and annotated in Jalview. **B:** Prediction of secondary structure in linker region of HsPINK1 and PhcPINK1 by SYMPRED. SYMPRED predicted α-helix represents a consensus prediction of α-helix by different prediction tools; PROF, SSPRO, YASPIN, JNET and PSIPRED. Predicted α-helices are highlighted by cylinders. **C:** Complete structure of human PINK1 solved by alphafold (Alphafold ID: Q9BXM7) and color-coded according to regions. **D:** PINK1’s α-NTE domain interaction with CTE domain where α-NTE region primarily interacts with αK region of CTE. **E:** Conserved residues at the interface along with PD-associated residues are highlighted. The residues of α-NTE at the start of interface (E117) and end of α-helix (K135) are also highlighted. **F:** Alphafold predicted Phpink1(108-575) model (grey) superposed with the crystal structure of insect pink1, Phpink1 (6EQI, blue) with a rmsd of 0.7. The missing NTE region in the crystal structure is labelled. **G:** HDX data mapped on Alphafold modelled *Ph*pink1^108-575^ with zoomed in view of NTE and CTE. The % deuterium uptake across the protein after 2min of labelling is color-coded as labelled. The residues are also color-coded according to the % deuterium uptake.

Furthermore, inspection of the human PINK1 AlphaFold model suggests that the N-terminal α-helix forms an intramolecular interface with the αK helix of the CTE (Figures 3C-E). Of relevance is the location of residues mutated in PD including Cys125 and Gln126 within this interface as well as Glu117. The key residues in the αK helix mediating the interaction include Val528, Leu532 and Leu539 (Figures 3C-E) in which a PD case has been reported with homozygous Leu539Phe mutation.

The AlphaFold human PINK1 structure model and PhcPINK1 crystal structure resemble structural arrangements with the NFK3 pseudokinase family members, PEAK1 and Pragmin that contain helical regions on either side of the pseudokinase domain, known, as Split Helical Dimerisation (SHeD) (Supplementary Figure 4A) [26, 27]. The N-terminal region (SHeD1) forms a single 24-residue α-helix (termed αS) similar to the predicted NTE domain for human PINK1 and PhcPINK1 (Supplementary Figure 4B). Most strikingly, the SHeD2 region of PEAK1 is highly similar to the PINK1 CTE domain with four α-helices (termed αJ-M) and superimposition of the PhcPINK1 CTE and PEAK1 SHeD2 reveals a high degree of structural similarity (Supplementary Figures 4C-E). PEAK1 utilises the surface of the αK and αL helices for a cis-contact with the SHeD1 αS-helix via ionic and hydrogen bond interactions (Supplementary Figure 4F). This structurally resembles the AlphaFold predicted interface in PINK1 which is mediated by the arrangement of hydrophobic residues on the surface of the NTE domain and αK-helix and the conservation of these hydrophobic residues in PhcPINK1 suggests a similar interaction of the NTE domain with the CTE domain (Supplementary Figure 4G).

We next expressed and purified an N-terminal extended fragment of PhcPINK1 (residues 108-575) containing the predicted NTE domain (Supplementary Figure 5) and performed HDX mass spectrometry (HDX-MS) to gain molecular insights into the putative NTE domain of PINK. HDX-MS gives relative rate of exchange backbone amide hydrogen with deuterium based on the strength of hydrogen bonding and solvent accessibility in a well ordered protein. It also can be used to distinguish peptides that are in a protein core (low uptake over time) from those that are exposed (high uptake) [28]. The HDX pepsin digestion yielded 185 peptides covering 85 % of the entire sequence (Supplementary Figure 6) and regions not covered were excluded from the analysis. Deuterium uptake after 2 min was used for the analysis as saturation was reached at this time. The rate of uptake was then colour coded and mapped on to an AlphaFold predicted structure of PhcPINK1 (residues 108-575) (Figures 3F-G). Peptides with uptake of less than 60% are considered not to be a random coil but a well-structured region. Consistent with the AlphaFold predicted model, the HDX analysis demonstrated that peptides within the NTE domain-containing region of PhcPINK1 exhibited low-medium level of deuterium uptake indicating that the region is well ordered (Figure 3G).

## Biochemical analysis indicates critical role of NTE:CTE interface for PINK1 activation in cells

The location of PD associated pathogenic mutations within the NTE domain suggests that the AlphaFold predicted NTE:CTE interaction is critical for PINK1 function. The generated model indicates that mutations in the conserved Cys125 residue (C125G) could disrupt the NTE:CTE interface whilst the Q126P mutation would break the NTE domain α-helix (Figure 3E). To investigate the functional impact of PD mutations at the interface, we generated stable cell lines in which we re-introduced full-length PINK1-3FLAG wild-type (WT); kinase-inactive mutant PINK1 (KI); NTE domain PINK1 mutants namely C125G, Q126P; CTE mutants PINK1 534_535InsQ and L539F, and kinase domain mutants namely A168P, E240K, G309D and G409V, into Flp-In™ T-REx™ Hela PINK1-knockout cells (generated by exon 2-targeted CRISPR-Cas9 (Supplementary Figure 7A-B)). Structural data from the insect orthologues indicate that A168P and E240K mutations disrupt ATP binding whilst G309D lies within INS3 and disrupts substrate binding without affecting autophosphorylation of Ser228 [9, 10]. G409V lies on the P+1 loop thereby preventing substrate recognition from the C-lobe side and has previously been reported to preserve autophosphorylation but disrupt downstream signalling [12].

To determine the effect of the selected PINK1 mutants on activation, cells were treated with DMSO or AO for 3 h to induce mitochondrial depolarisation. Mitochondrial-enriched fractions were isolated and solubilized in 1% Triton-X100 lysis buffer. Immunoblot analysis of PINK1 demonstrated significant reduction of PINK1 levels in mitochondria for the N-terminal mutants C125G, Q126P and the CTE mutant 534_535InsQ compared to WT PINK1 following mitochondrial depolarisation (Figures 4A-B). We also observed decreased PINK1 expression in the ATP binding defective mutants, A168P and E240K, but there was no alteration of expression for the substrate binding-defective mutants, G309D and G409V (Figures 4A-B). To monitor autophosphorylation, we employed a polyclonal phospho-specific Ser228 antibody (Waddell et al, manuscript in preparation) (see Materials and Methods) and observed complete loss of Ser228 autophosphorylation in the C125G, Q126P, and 534_535InsQ mutants similar to PINK1 KI expressing cells (Figures 4A-B). We observed no reduction of Ser228 autophosphorylation in the substrate binding mutants, G309D and G409V, which is consistent with previous studies of these mutants in insect PINK1 (G309D) and mammalian cells (G409V) [9, 10, 12, 29]. Interestingly we saw residual Ser228 autophosphorylation in the A168P and E240K mutant suggesting that these do not completely abolish catalytic activity (Figures 4A-B). Finally C125G, Q126P, A168P, E240K, G309D, G409V and 534_535InsQ were associated with complete loss of phospho-ubiquitin upon mitochondrial depolarisation, that would disrupt downstream signalling consistent with their pathogenic role in the development of Parkinson’s disease (PD) (Figures 4A-B).

**Figure 4.**
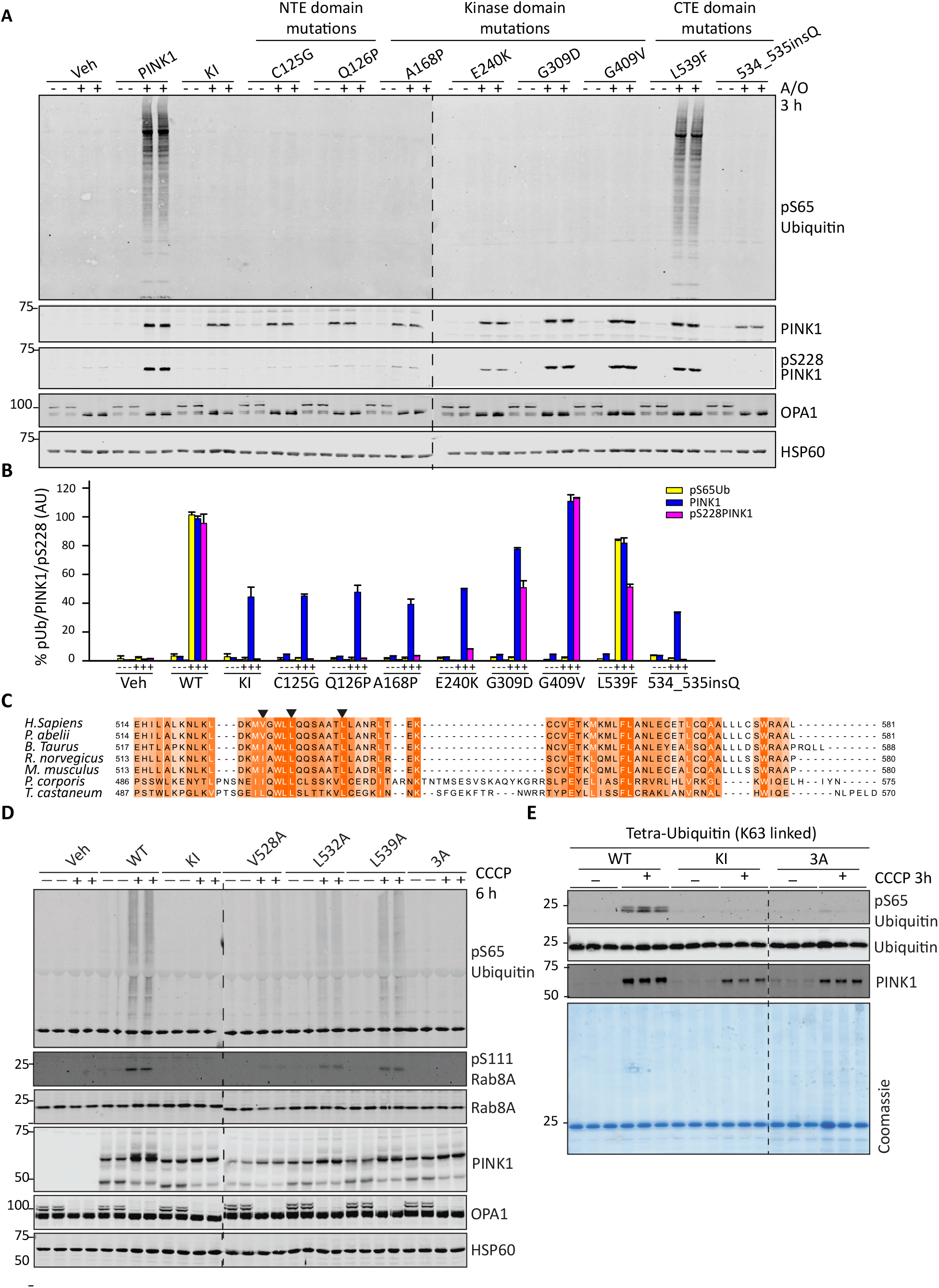
Mutational analysis confirms critical role of NTE and CTE domains for PINK1 activation. **A:** α-NTE mutants and CTE PD-associated mutants results in loss of autophosphorylation. Stably-expressing PINK1-3FLAG WT, KI (D384A), empty, α-NTE mutants (C125G, Q126P), kinase domain mutants (A168P, E240K, G309D, G409V) and CTE domain mutants (L539F, ins534Q) cell lines were generated in PINK1 knockout Hela Flip-in cells. PINK1-3FLAG expression was induced by 24 hr treatment with 0.2 uM doxycycline, with mitochondrial depolarized by 3 hr treatment with 10µM A/O where indicated. Mitochondria-enriched fractions were subjected to immunoblotting with α-PINK1 (DCP), α-Ubiquitin pS65 (21^st^ Century), α-OPA1 (BD) and α-HSP-60 primary antibodies. N=3. **B:** Immunoblots were quantified for phospho-Ser65 Ub/HSP-60, PINK1/HSP-60 and pS228/HSP- 60 using the Image Studio software. Data are presented relative to the PINK1-WT as mean ± SD (n=3). **C:** Multiple sequence alignment of CTE region of PINK1 orthologs across species. Sequence alignment was performed with MUSCLE and annotated in Jalview. Mutated CTE residues for functional analysis are highlighted with arrow heads. **D:** C-terminal extension (CTE) mutants exhibit reduced stabilisation, auto-phosphorylation and substrate phosphorylation. hPINK1 knock out HeLa cells transiently expressing 3xFLAG tagged hPINK1 WT, KI or hPINK1 CTE mutants V528A (L504A in TcPINK1), L532A (L508A) and L539A (L515A). Cells were stimulated -/+ 10 μM CCCP for 6 hr. Membrane fractions were isolated and solubilised in 1% tritonx100 lysis buffer Lysates were resolved by SDS PAGE. Proteins were transferred to nitrocellulose membranes probed using the antibodies indicated. n=2. **E:** CTE triple mutant is inactive against recombinant substrates in vitro. Flp-In HEK293 PINK1 KO cells stably expressing WT or KI (D384A) or dimerization triple mutant (V528A/L532A/L539A) PINK1 were treated -/+ 10µM CCCP for 3 hours and subjected to sub cellular fraction in a cell free assay buffer (CFAB). 3µg of non-solubilised membrane enriched fraction (MeF) were incubated with tetraubiquitin (K63 linked) in CFAB supplemented with 2mM (unlabelled) ATP, 5mM MgCl2, 2mM DTT and 0.75% Glycerol for 20mins. Samples were resolved via SDS page and blotted for the indicated antibodies. N=2

Under these assay conditions we did not observe any impact of the L539F mutation on PINK1 stabilisation, autophosphorylation or ubiquitin phosphorylation (Figures 4A-B). Based on the AlphaFold structure model, the NTE:CTE interface is facilitated by key hydrophobic residues on the surface of the αK-helix composed of Val528, Leu532 and Leu539 residues that are well-conserved across species (Figures 3E and 4C). Since, Phenylalanine (Phe) is also hydrophobic, it would be anticipated that the L539F mutation would not be deleterious to the interface. To validate the functional role of the hydrophobic CTE interface, we transiently transfected WT and KI PINK1-3FLAG alongside CTE mutants, V528A, L532A, L539A and a combinatorial V528A/L532A/L539A triple mutant (3A) in Flp-In™ T-REx™ HeLa PINK1-knockout cells followed by 6 h treatment with DMSO or 10 μM CCCP. Immunoblot analysis of PINK1 in mitochondrial enriched fractions demonstrated reduced PINK1 levels for all the CTE mutants, V528A, L532A, L539A, as well as in the 3A mutant compared to controls (Figure 4D). Activation of PINK1 was assessed by immunoblotting against substrates, phospho-ubiquitin and phospho-Ser111-Rab8A (pSerRab8A) and all mutants led to reduced phospho-ubiquitin and pSerRab8A – most notable in the V528A, L532A mutants – and a lesser reduction in the L539A mutant (Figure 4D). Strikingly, PINK1 activation was completely abolished in the 3A mutant (Figure 4D). To further assess how the CTE mutants affect mitochondrial localization and activity of PINK1, we performed an *in vitro* intact mitochondrial kinase assay of PINK1 (Supplementary Figure 8) [18]. We isolated mitochondria-enriched fractions from Flp-In™ T-REx™ HEK293 PINK1-knockout cells stably re-expressing WT, KI or the 3A mutant of human PINK1and subjected these extracts to *in vitro* kinase assay using recombinant K63-linked tetraubiquitin (K63-Ub4) (Figure 4E). Immunoblotting of PINK1 demonstrated decreased PINK1 levels in the isolated mitochondria of PINK1 KI and 3A expressing cells following mitochondrial depolarisation (Figure 4E). Under these assay conditions, WT PINK1 that was localised to mitochondria efficiently phosphorylates K63-Ub4, however, this was completely abolished in mitochondria isolated from KI PINK1 and 3A PINK1-expressing cells (Figure 4E). Overall these studies demonstrate the critical role of the CTE hydrophobic residues in PINK1 activation and strongly suggest that the L539F mutant is not pathogenic and causal of PD in the case reported [30].

The location of the NTE:CTE interface outside the kinase domain suggests that mutants would not directly affect the intrinsic catalytic activity of PINK1. Previous studies have found that expression of recombinant human PINK1 in *E*. coli displays very low catalytic activity whereas expression of insect PINK1 orthologues exhibit robust activity *in vitro*. To determine the impact of NTE domain and CTE mutations on PINK1 catalytic activity, we performed an *in vitro* kinase using recombinant WT and mutant PhcPINK1 (residues 108-575) expressed in *E. coli*, incubated with [γ-^32^P]-ATP and K63-Ub4 substrate. Consistent with the AlphaFold structure prediction, we observed that all mutants tested namely, E125K, C128G, W129P, I503A, L507A, L514F (equivalent to human E117K, C125G, Q126P, V528A, L532A, L539F respectively) efficiently phosphorylated K63-Ub4, similar to WT PINK1 as determined by measuring [γ-^32^P]-ATP incorporation via autoradiography (Supplementary Figure 9A). To determine whether NTE domain mutants affect the ability of PINK1 to associate with the TOM complex we generated mitochondrial-enriched fractions from HeLa PINK1 KO cell lines stably re-expressing WT, KI, S228A, C125G and Q126P mutant PINK1, and subjected these to BN-PAGE assays. Consistent with previous studies we observed that WT PINK1 and the S228A PINK1 mutant stabilise in a ∼700kDa complex in response to mitochondrial depolarisation (Supplementary Figure 9B). We observed that the C125G and Q126P had reduced PINK1 levels at the mitochondria similar to KI PINK1, but there was striking loss of association with the TOM complex to a greater degree than KI PINK1 (Supplementary Figure 9B). This suggests that a major role of the NTE:CTE interface is to facilitate PINK1 stabilisation at the TOM complex following mitochondrial depolarisation.

### Decreased stabilisation and activation of endogenous PINK1 in homozygous *PINK1*^*Q126P*^ patient-derived fibroblasts

We previously identified two sisters carrying homozygous mutations in PINK1, p.Q126P [15]. Both sisters manifested with early onset PD at the age of 37 and 41 years and were longitudinally assessed in the dopaminergic ON state at the outpatient clinic for Parkinson’s Disease, University Hospital Tuebingen. At the most recent follow-up, the disease duration was 39 and 30 years, respectively. Both sisters had only mild motor impairment (UPDRS-III: 15 and 20 points after 27 and 30 years disease duration) with good Levodopa response. Cognitive function was mildly impaired with 21 and 23 points in MoCA testing after 27 and 30 years disease duration. No manifest mood disturbances were seen (BDI 11 and 10 points after 27 and 30 years disease duration). In line with the benign disease course, CSF levels of Amyloid-beta1_42 (729 and 678 pg/ml; cut-off ≥600pg/ml), total-Tau (159 and 196 pg/ml; cut-off ≤450pg/ml), phosphorylated-Tau (40 and 37 pg/ml; cut-off ≤60pg/ml) and NFL (640 and 598 pg/ml; cut-off ≤966pg/ml) were normal after 26 and 29 years disease duration. Both sister showed no α-synuclein seeding activity using RT-QuIC assay. Primary skin fibroblast cultures of one sister with homozygous PINK1 Q126P mutation, and unaffected human control fibroblasts, were treated with 10 μM CCCP or DMSO for 3 h or 24 h (Figure 5). Immunoprecipitation-Immunoblotting with anti-PINK1 antibody demonstrated time-dependent stabilisation of PINK1 in control fibroblasts but this was significantly reduced in fibroblasts expressing Q126P PINK1 (Figure 5). This was associated with complete loss of phospho-ubiquitin and pSer111Rab8A in Q126P fibroblasts compared to control cells, indicating reduced PINK1 activation (Figure 5).

**Figure 5.**
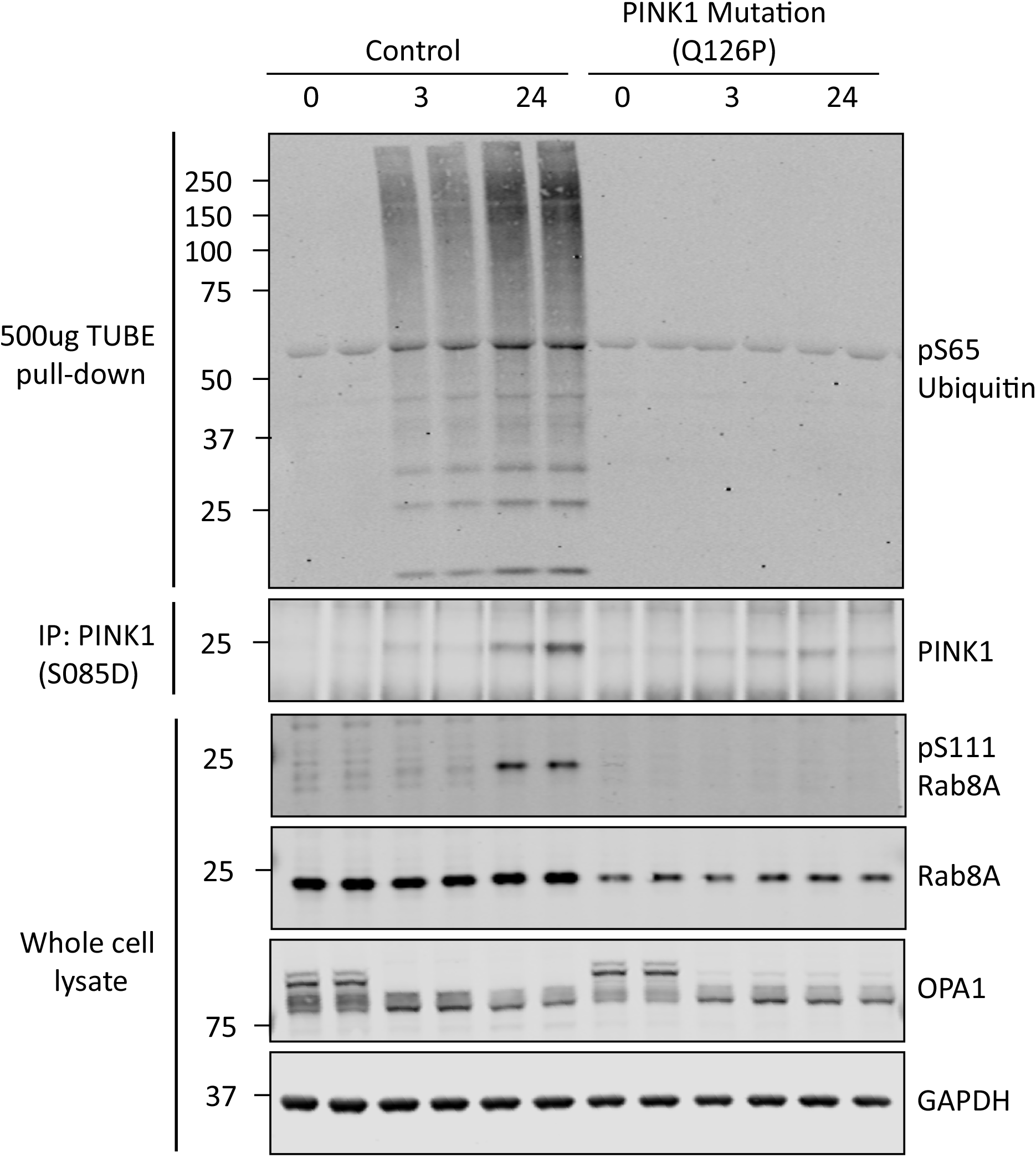
Endogenous PINK1 stabilisation and activation is reduced in Q126P PINK1 patient derived fibroblasts. Primary fibroblast cultures established from skin biopsies from a healthy subject (Control) and a PD patient harbouring a PINK1 Q126P homozygous mutation were treated with 10 μM CCCP or DMSO for 3 h or 24 h. Lysates were subjected to immunoblot analysis using the indicated antibodies. Ubiquitin-capture was used prior to immunoblotting with anti-phospho-Ser65 ubiquitin antibody. Endogenous PINK1 was detected after immunoprecipitation from whole cell lysate.

## Discussion

Combining mutational analysis with bioinformatic structural predictions of human PINK1, we have been able to map and elaborate a functional role for an N-terminal α-helical extension (NTE domain) of human PINK1 (Figures 2, 3 and Supplementary Figure 2). We demonstrate a critical role of the interaction of the NTE domain with the CTE domain, towards PINK1 stabilisation, Ser228 autophosphorylation and activation, and that human disease-causing mutations of PINK1 that lie within the NTE-CTE interface disrupt PINK1 activation (Figure 4 and Supplementary Figure 9). Finally we report the impact of homozygous NTE mutation, Q126P, on preventing stabilisation and activation of endogenous PINK1 in patient-derived fibroblasts (Figure 5).

PINK1 stabilisation is regulated by multiple factors, notably the abrogation of proteolysis by PARL and other proteases upon mitochondrial depolarisation [31]. Active PINK1 is stabilised in the Translocase of outer membrane (TOM) complex [11, 12], however, to date the molecular mechanism of how this contributes to PINK1 activation remain to be fully elucidated. Interestingly we observed that the HSP60 mitochondrial targeting sequence (MTS) (residues 1-27) and PINK1 MTS (residues 1-34) both permit PINK1 import however, the PINK1 MTS did not lead to PINK1 activation. In future work it will be interesting to understand whether the mitochondrial import mechanism of PINK1 MTS and HSP60 MTS-PINK1 is different, or whether they are both imported via the TOM complex but their association with it upon mitochondrial depolarisation, is distinct.

Previous analysis of PINK1 mutants have found that disease mutants that affect intrinsic catalytic activity disrupt PINK1 stabilisation at the TOM complex and this is consistent with our observations with kinase-inactive and ATP-binding defective mutants of PINK1 (Figure 4) [11, 12]. The finding that NTE or CTE domain mutants have similar impacts on PINK1 stabilisation at the TOM complex independent of intrinsic catalytic activity provide new insights into PINK1 stabilisation (Figures 4, 5 and Supplementary Figure 89?). Studies in TOM7 knockout cells have demonstrated that TOM7 is essential for PINK1 kinase activation [32]. A recently uploaded preprint has described the structure of *Tribolium castaneum* PINK1 (TcPINK1) that provides direct experimental evidence for the NTE:CTE molecular interaction [33]. Future structural studies will be required to determine whether the NTE:CTE interface facilitates PINK1 binding directly to TOM7

Interestingly we observed that substrate binding-defective mutants, G309D and G409V, that exhibit robust autophosphorylation at Ser228, stabilise as well as wild-type PINK1 (Figure 4). This suggests that autophosphorylation of PINK1 rather than substrate phosphorylation of PINK1 contributes to PINK1 stabilisation. We found that the S228A mutant was still able to stabilise at the TOM complex (Supplementary Figure 9B), in agreement with a previous study [12], and the identity of PINK1 autophosphorylation sites mediating PINK1 stabilisation at the TOM complex remain to be discovered.

We did not observe any biochemical defect of the L539F mutation on PINK1 activation (Figure 4). This mutation has been reported in only one patient/family with PD compared to the other mutations we assessed which have been found in multiple independent families, and overall our data indicates that this mutation is not pathogenic and unlikely to be the cause of Parkinson’s in the reported case [30]. In contrast the clinical relevance and importance of the function of the NTE:CTE interface is supported by the demonstration of defective endogenous PINK1 stabilisation and activation in patient-derived fibroblasts of a homozygous Q126P mutation. The age of onset of the patient was < 45 years and the slow progression is clinically consistent with PINK1 pathogenic mutations in humans. Identifying further patients harbouring pathogenic mutations at the NTE:CTE interface will be important to aid in translational studies exploiting this regulatory mechanism of PINK1.

In conclusion our studies have revealed an important functional role for the NTE α-helical domain of PINK1 and its interaction with the CTE as a mechanism required for PINK1 stabilisation at the TOM complex (Figure 6). Our findings suggest that small molecules that target the NTE:CTE interface to enhance association with the TOM complex could have therapeutic benefit in select PINK1 patients, and the feasibility of this approach will be aided by further structural analysis.

**Figure 6.**
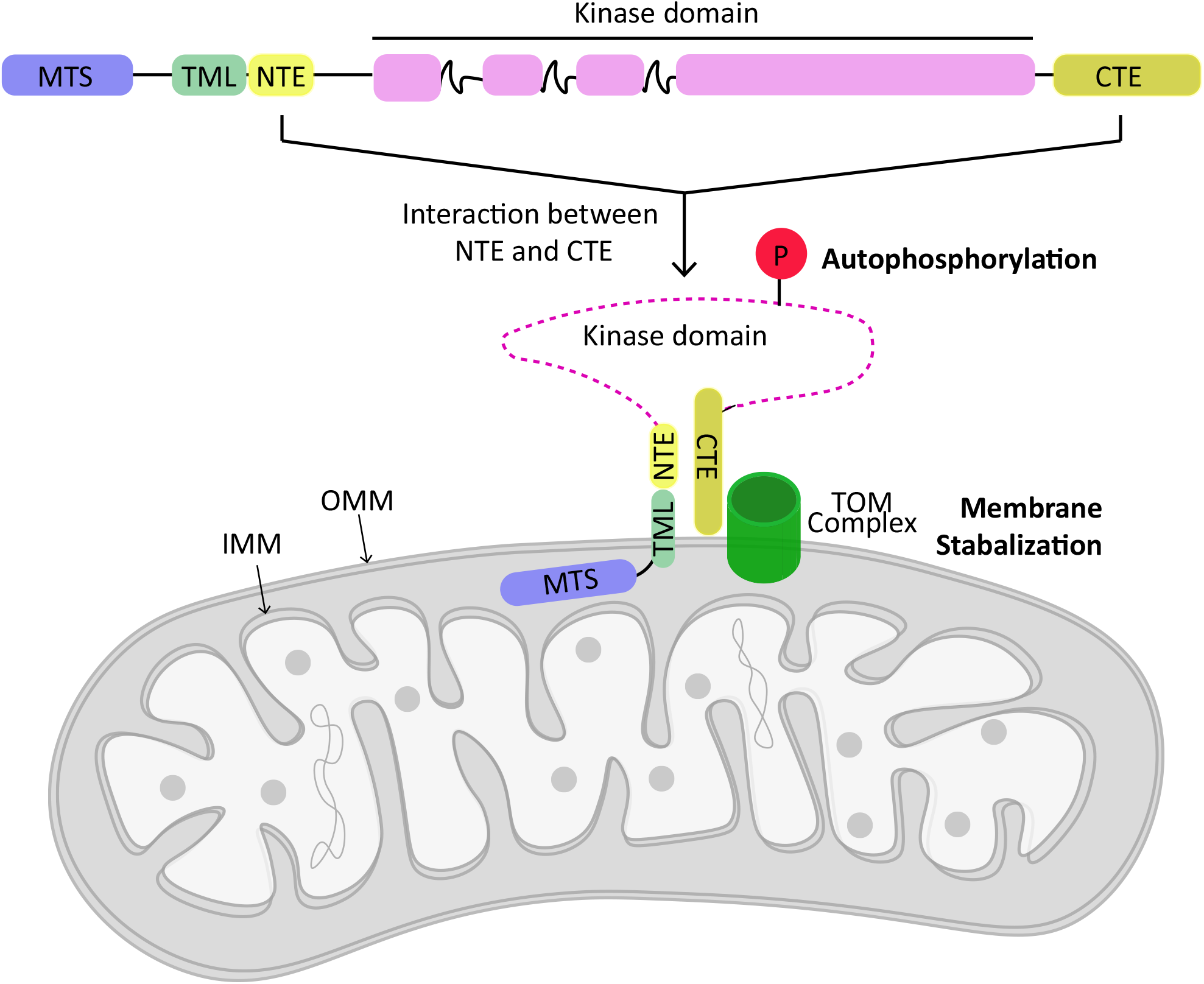
Schematic of role of NTE:CTE interaction towards PINK1 stabilisation at mitochondria. NTE domain interaction with CTE facilitates stabilisation and activation of PINK1 at TOM complex. Abbreviations: mitochondrial targeting sequence (MTS); transmembrane-like domain (TML); N-terminal extension domain (NTE); C-terminal extension (CTE); Translocase of outer membrane (TOM); inner mitochondrial membrane (IMM); outer mitochondrial membrane (OMM).

## Acknowledgements

We thank Daan van Aalten and Yogesh Kulathu for helpful discussions. We express our thanks to Mark Peggie, Simone Weidlich, Mel Wightman and Rachel Toth for molecular biology and cloning. This work was supported by a Wellcome Trust Senior Research Fellowship in Clinical Science (210753/Z/18/Z to M.M.K.M); the Michael J. Fox Foundation (R.B. and M.M.K.M.), the Medical Research Council [MR/P00704X/1 COEN award] (M.M.K.M); EMBO YIP Award (M.M.K.M.); CRUK Programme Award (C24461/A23303 to R.B.); and the German Federal Ministry of Education and Research (BMBF; PDStrat; FKZ 031L0137B) in the frame of ERACoSysMed2 (K.B.). A.S. was recipient of a BBSRC EASTBIO PhD Studentship. E.M.K. was recipient of an Astex Pharmaceuticals Sustaining Innovation Post-Doctoral Award. We are grateful to the sequencing service (School of Life Sciences, University of Dundee); Axel Knebel for expression and generation of K63-tetra-ubiquitin and TUBE proteins (MRC PPU); the MRC PPU tissue culture team (co-ordinated by Edwin Allen) and MRC PPU Reagents and Services antibody teams (co-ordinated by James Hastie).

## Author Contributions

PK, HO, OR, AS, ADW, and SB performed experiments and analysed data. AK, EMK, MSA contributed to biochemical analysis. JRA performed HDX-MS; KB and JCF isolated and characterized human fibroblasts and generated clinical data; TM designed and generated CRISPR constructs. MMKM and RB supervised experiments and analysed results. PK, HO, and MMKM wrote the paper with contribution from all the authors. MMKM conceived and supervised the project.

## Conflict of Interest

M.M.K.M. is a member of the Scientific Advisory Board of Mitokinin Inc.

**Supplementary Figure 1. Generation of Flp-In™ T-REx™ HEK293 PINK1-knockout cell lines by CRISPR-Cas9. A:** Genetic sequencing of Flp-In™ T-REx™ HEK293 cell line clone “F1” following exon 2-targeted CRISPR/cas9. Wildtype PINK1 sequence is indicated (PINK1). Knockout clone sequencing is indicating for both Allele 1 and Allele 2. Both allele’s possess frameshift mutations starting at residue Leu143 with resulting stop codons following 22 or 2 amino acids, respectively. **B:** Immunoblot characterisation of WT (293) and PINK1-knockout (293 k/o) HEK293 cells. Indicated cell lines were treated for 3 hr with 10 μM CCCP, with samples then analysed for PINK1 expression by immunoblot.

**Supplementary Figure 2. Structural modelling of N-terminal α-helix extension (NTE domain) of human PINK1. A:** Schematic representation of human PINK1 (HsPINK1) and Pediculus humanis corporis PINK1 (PhcPINK1) highlighting the linker region flanked by transmembrane domain and kinase domain. The alignment of linker region is performed by MAFT and annotated in Jalview. **B, C:** Detailed prediction of secondary structure in linker region of HsPINK1 and PhcPINK1 by SYMPRED. SYMPRED predicted α-helix represents a consensus prediction of α-helix by different prediction tools; PROF, SSPRO, YASPIN, JNET and PSIPRED. Predicted α-helices are highlighted by cylinders. **D:** Human PINK1 structure predicted by Alphafold (Alphafold ID: Q9BXM7) is color-coded according to domains arrangement (left) and model confidence score, pLDDT (right). Zoomed in view of NTE and CTE showed high model confidence score.

**Supplementary Figure 3. Human PINK1 alignment with PINK1 orthologues**. Alignment performed with MUSCLE and annotated with Jalview. The mitochondrial targeting sequence (MTS) is highlighted by a blue box and the transmembrane-like domain (TML) is highlighted by green box. The predicted N-terminal extension (NTE domain) is highlighted by a yellow cylinder. The kinase domain is highlighted with pink box, in which first, second and third insertions are highlighted with loops. Amino acid motifs critical for catalysis are highlighted as follows – yellow: glycine rich loop; orange: LAIK motif lysine; dark red: HRD catalytic motif; purple: DFG motif/N-terminal activation loop anchor; green: APE motif/C-terminal activation loop anchor. The CTE domain is highlighted with magenta box.

**Supplementary Figure 4. Structural modelling of interaction of PhcPINK1 NT and CTE domain based on PEAK1 SHeD1 and SHeD2 domains. A:** Schematic representation of PhcPINK1 and PEAK1. Dotted lines indicate the possible similarity of CTE domain with SHeD2 domain and α-NTE domain with SHeD1 domain. **B:** Sequence similarity between α-helix region of HsPINK1, PhcPINK1 and PEAK1/Pragmin SheD1 domain. Sequence alignment were performed using MAFT and annotated by Jalview. The asterisks indicate PD associated mutant positions in human lying across a fairly conserved region. **C:** Structural similarity between PhcPINK1 and NF3 Pseudokinase PEAK1. Structural alignment of CTE domain and SHeD2 domain is performed by DALI and annotated by Jalview. Unsolved loops are represented by hashes and α-helices (a-J,K,L,M) of SHeD2 domain and CTE domain are shown as cylinders. **D:** Structure of PEAK1 SHeD2 domain (PDB ID: 6BHC) and PhcPINK CTE domain (PDB ID: 6EQI) are shown according to the solved crystal structures. **E:** Structural superimposition of PhcPINK1 CTE domain and PEAK1 SHeD2 domain is performed in SALIGN. **F:** Interaction of PEAK1’s SHeD1 and SHeD2 domains as shown in solved crystal structure (PDB ID: 6BHC). The αS of SHeD1 primarily interacts with αK and αL helices of SHeD2. The hydrophobic patches at the interface are shown as yellow bubbles. **G:** Hydrophobic residues are present at αK and αL of CTE domain of PhcPINK1 (PDB ID: 6EQI). Hydrophobic residues, primarily I503 and L507 are facing outwards in αK helix suggesting a potential interface. A suggestive position for conserved L514 residue position is shown by dotted line (unsolved structure). Model of predicted PhcPINK1 α-helix (made by SWISS-MODEL) shows hydrophobic patch comparable to SheD1 domain. A suggestive mode of binding between PhcPINK1 α-helix and CTE domain facilitated by hydrophobic regions of CTE and PhcPINK’s predicted α-helix is indicated by arrow.

**Supplementary Figure 5. Expression and purification of Pediculus PINK1. A:** SDS-PAGE profile of Phc PINK1 (108-575). **B:** Protein was purified on a superdex75 60/600 column, flow rate 1.0 ml/min. The elution profile reveals a small overlapping oligomeric state of the protein with the majorly monomeric profile.

**Supplementary Figure 6. Sequence coverage of HDX-MS mapping of Pediculus PINK1**. HDX-MS experiment sequence coverage for wild-type Phc PINK1 (108-575) at 10 µM deuterium labelling. A total of 122 peptides were identified amounting to 70% coverage. Peptides shown are those that pass the identification criteria as described in the method section. Residues are numbered as for the full-length protein.

**Supplementary Figure 7. Generation of Flp-In™ T-REx™ Hela PINK1-knockout cell lines by CRISPR-Cas9. A:** Genetic sequencing of Flp-In™ T-REx™ HeLa cell line clone “3C9” following exon 2-targeted CRISPR/cas9. Wildtype PINK1 sequence is indicated (PINK1). Knockout clone sequencing is indicating for both Allele 1 and Allele 2. Allels 1 and 2 possess frameshift mutations starting at residue Leu143 and Pro140 with resulting stop codons following 13 or 8 amino acids, respectively. **B:** Immunoblot characterisation of PINK1-knockout (Hela k/o) Hela cells in comparison to k/o cells with transfected PINK-GFP. Indicated cell lines were treated for 9 hr with 10 μM A/O, with samples then analysed for PINK1 expression by immunoblot.

**Supplementary Figure 8. Intact mitochondrial human PINK1 kinase assay. A:** Intact mitochondrial kinase assay (MitoKA) against tetra-ubiquitin (K63 linked) Flp-In TRex HEK293 PINK KO cells stably expressing WT or KI (D384A) PINK1 were treated -/+ CCCP for 3 hours and subjected to sub cellular fraction in a cell free assay buffer (CFAB). Indicated amounts of non-solubilised membrane enriched fraction (MEF) were incubated with 2µM tetra-ubiquitin (K63 linked) in CFAB supplemented with 2mM (unlabelled) ATP, 5mM MgCl2, 2mM DTT and 0.75% Glycerol for 1 hour. Samples were resolved via SDS page and blotted for the indicated antibodies. N=2 **B:** Time point optimisation for MitoKA against tetra-ubiquitin (K63 linked) Flp-In TRex HEK293 PINK KO cells stably expressing WT PINK1 were treated -/+ CCCP for 3 hours and subjected to sub cellular fraction in a cell free assay buffer (CFAB). Indicated amounts of non-solubilised membrane enriched fraction (MeF) were incubated with 2µM tetra-ubiquitin (K63 linked) in CFAB supplemented with 2mM (unlabelled) ATP, 5mM MgCl2, 2mM DTT and 0.75% Glycerol for indicated time points. Samples were resolved via SDS page and blotted for the indicated antibodies Red * indicates misloaded replicate. N=2

**Supplementary Figure 9. NTE domain and CTE domain mutations do not impact intrinsic catalytic activity of PhcPINK1. A:** Ubiquitin phosphorylation assay was performed with 250 nM PhcPINK1 (WT), kinase inactive (D357A; KI) and indicated mutations within the NTE and CTE domain and 2 µM K63-linked ubiquitin tetramer as substrate. Samples were subjected to SDS-PAGE, and analysed by Coomassie staining and [γ-^32^P] incorporation measured by autoradiography with Cerenkov counting. **B:** Mitochondria enriched fractions from Hela cells stably expressing PINK1-WT, KI, and PD-associated NTE domain mutants were treated with AO for 3 h and then subjected to BN-PAGE and immunoblotted for anti-FLAG. Samples were also subjected to SDS-PAGE and immunoblotting with anti-FLAG, OPA-1 and HSP-60 antibodies.

