## Supplementary Figures for "Mapping of a N-terminal α-helix domain required for human PINK1 stabilisation, Serine228 autophosphorylation and activation in cells"

**Supplementary Figure 1**

A

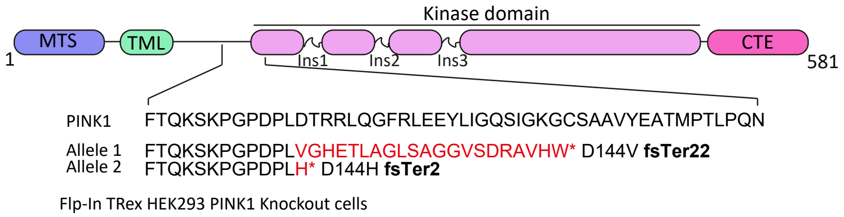


B

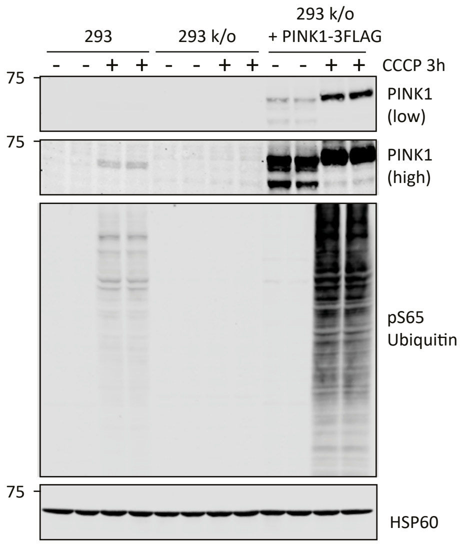

**Supplementary Figure 2**

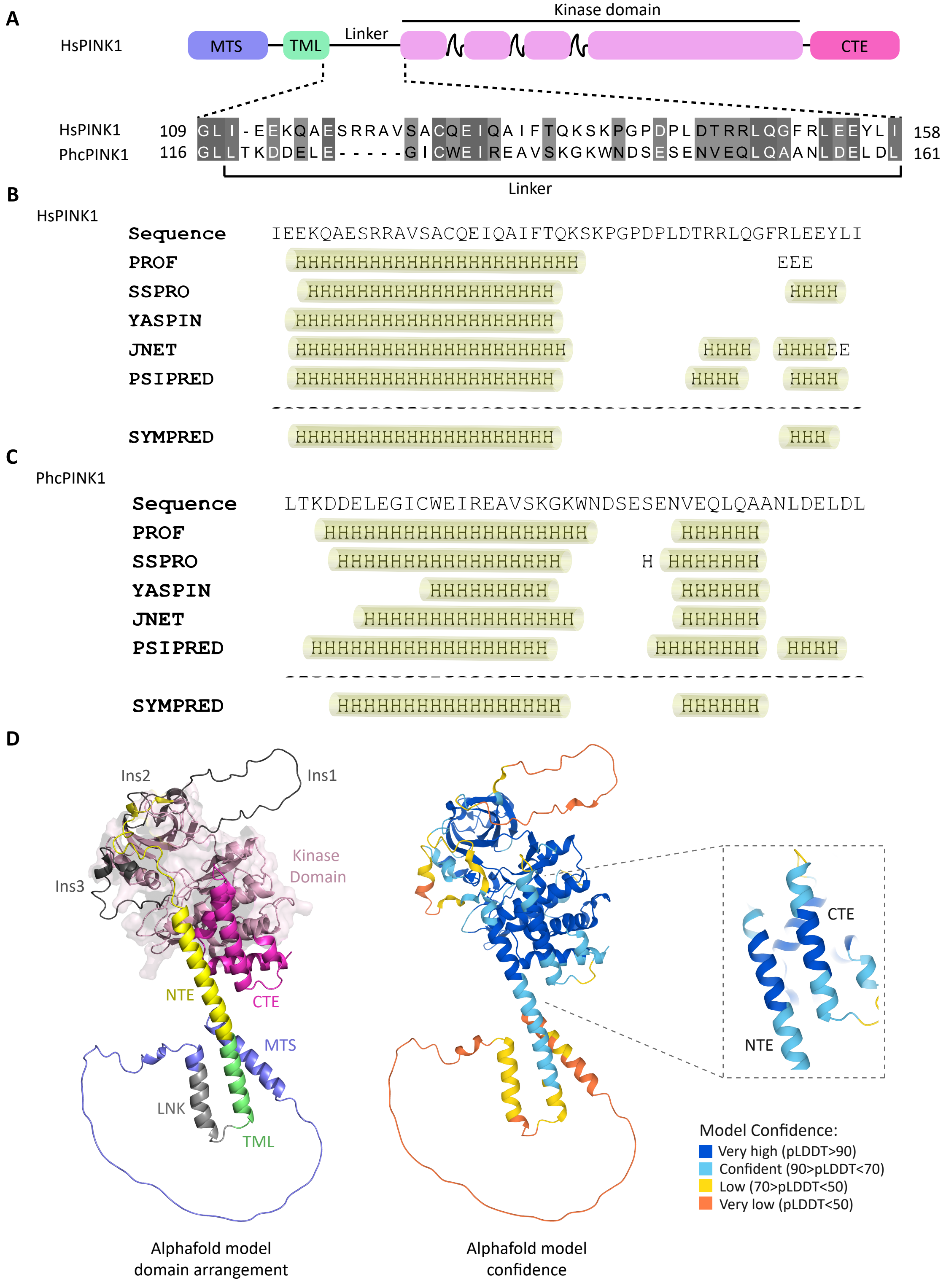

### Supplementary Figure 3

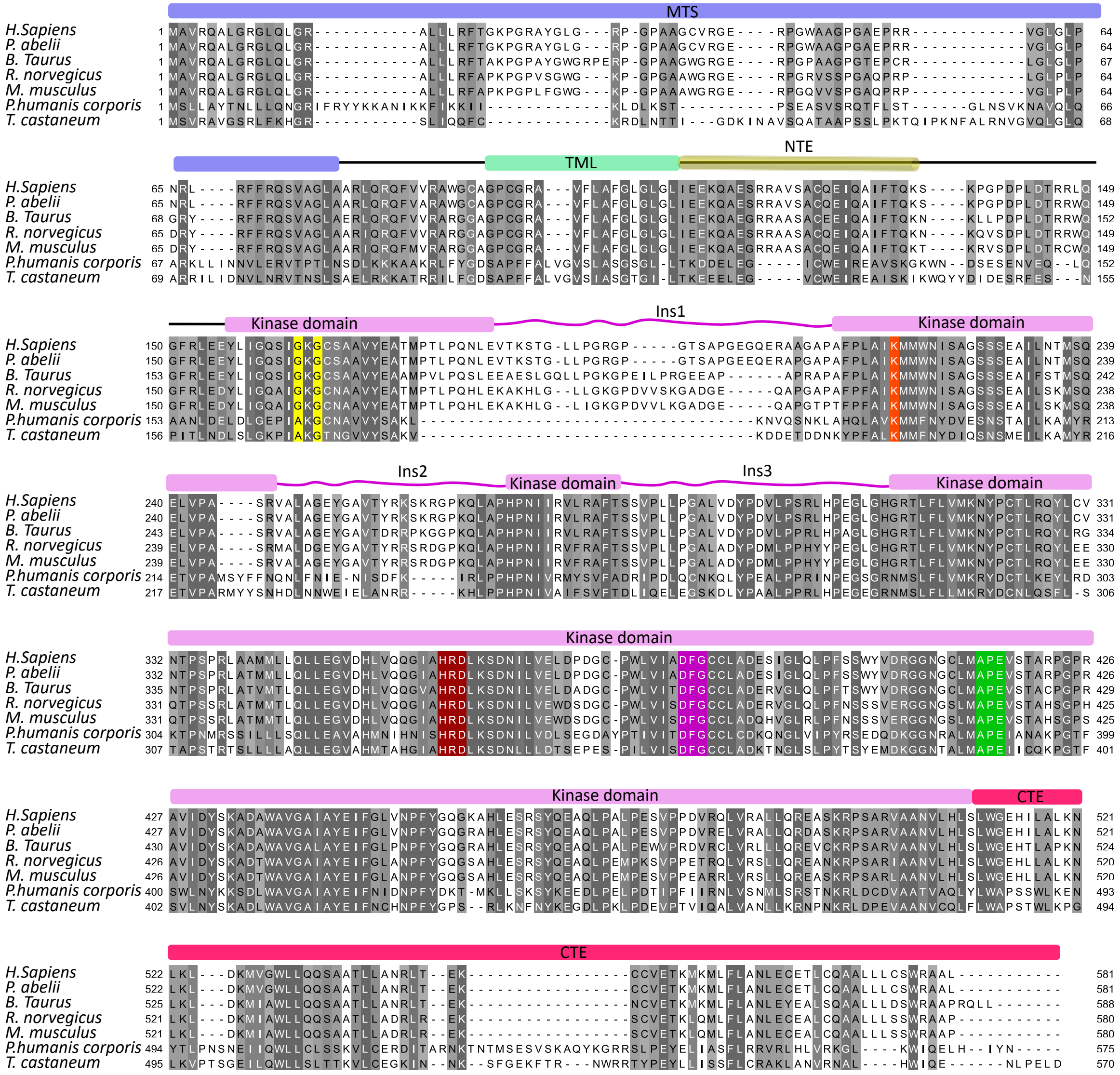

#### Supplementary Figure 4

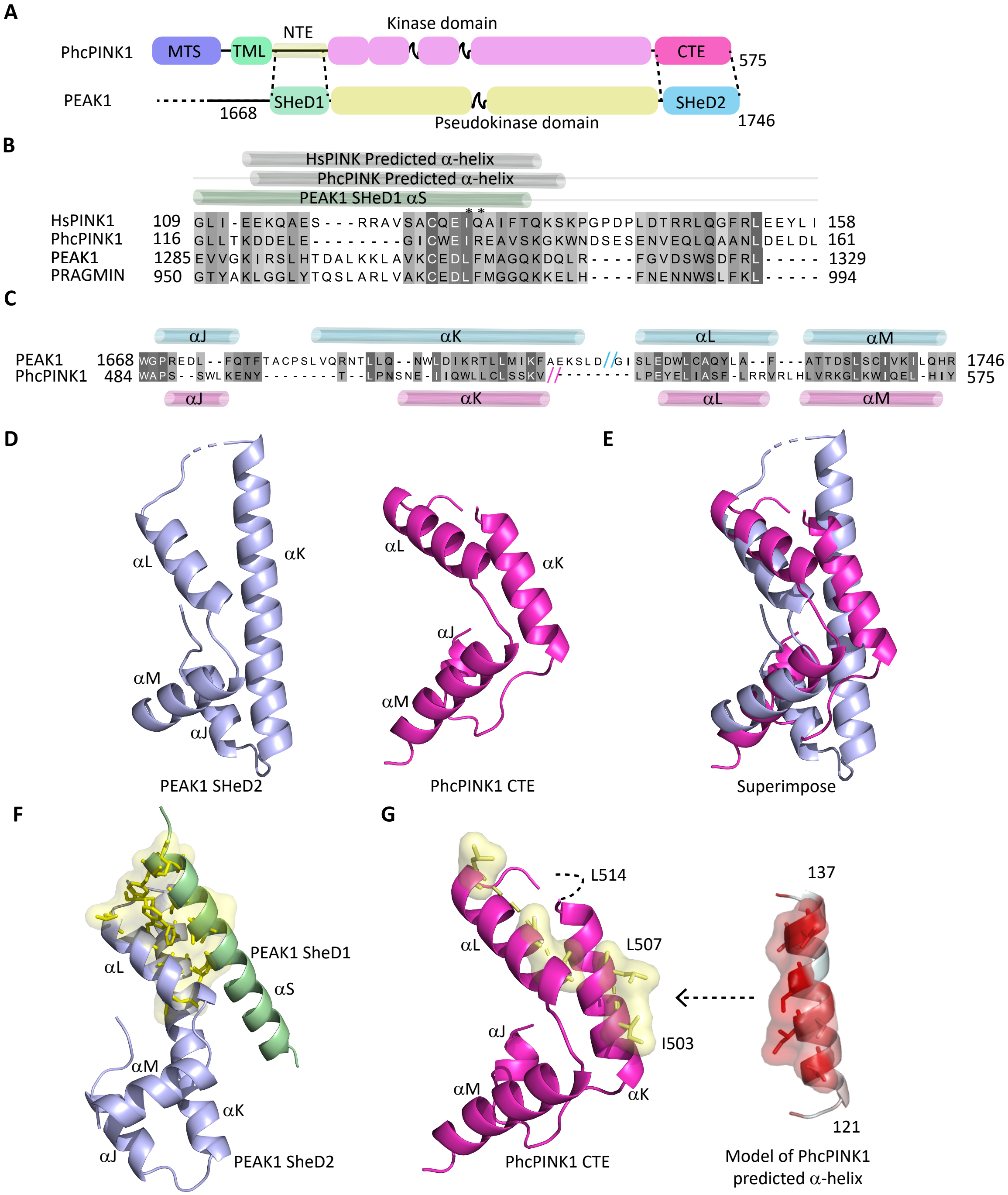

#### Supplementary Figure 6

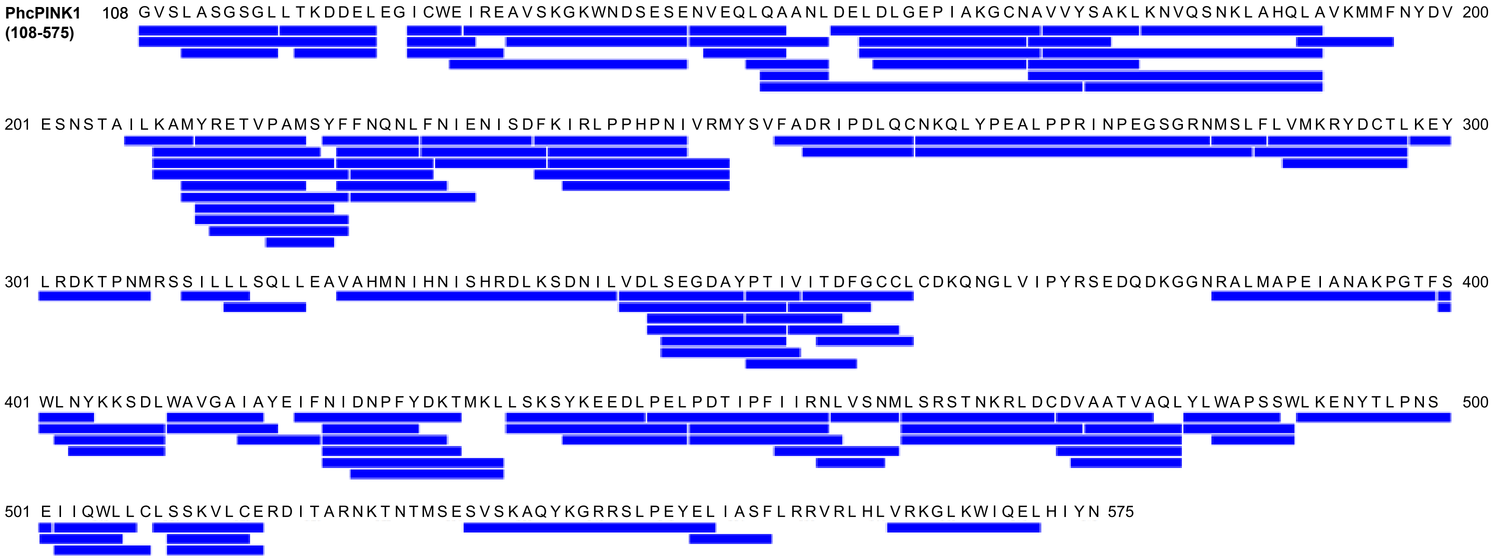

#### Supplementary Figure 5

**A**

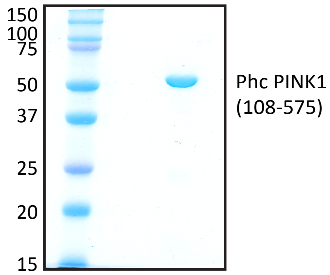


**B**

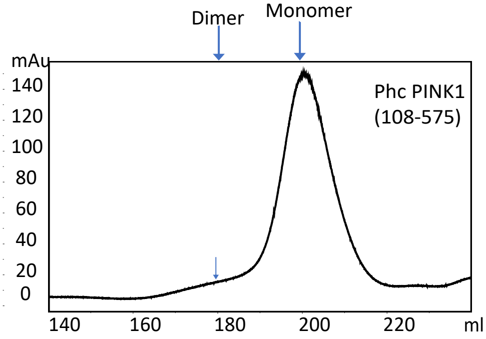

#### Supplementary Figure 7

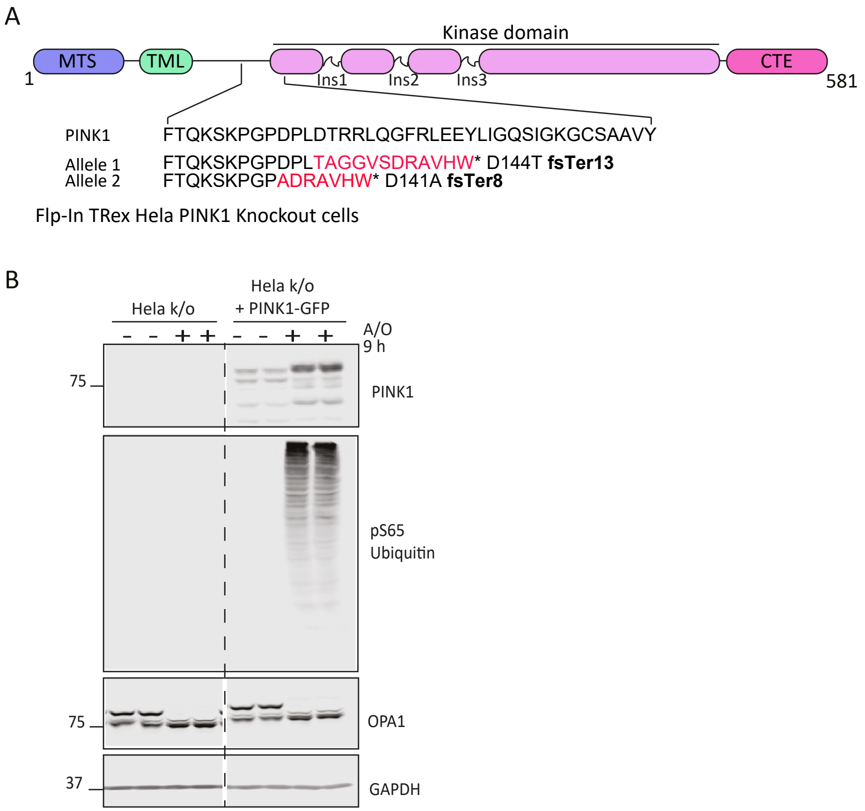

#### Supplementary Figure 8

**A**

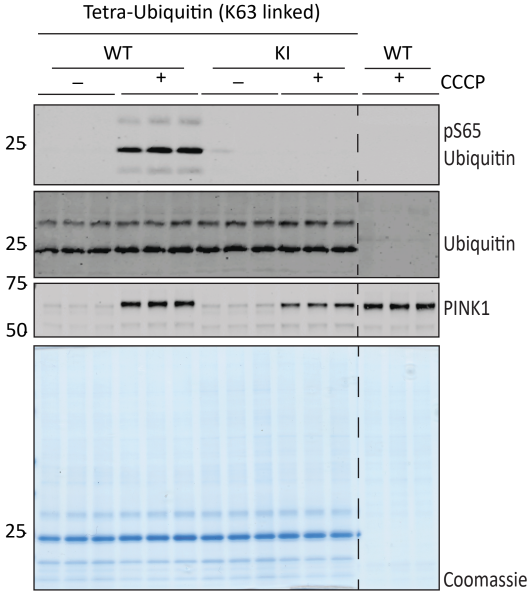


**B**

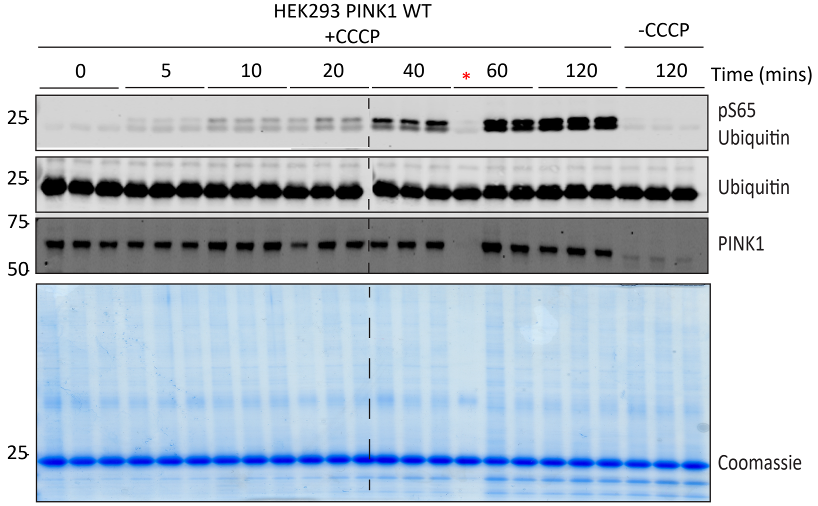

#### Supplementary Figure 9

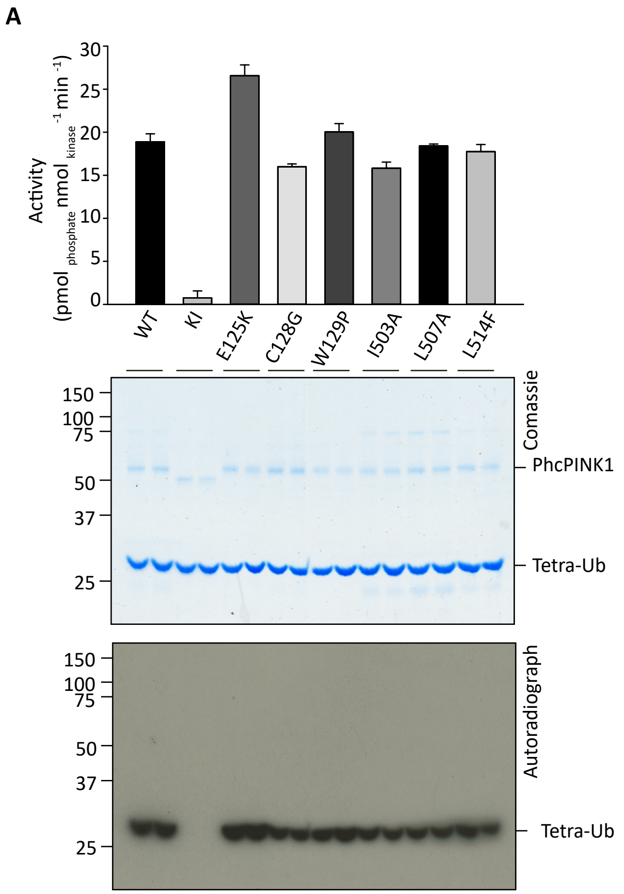


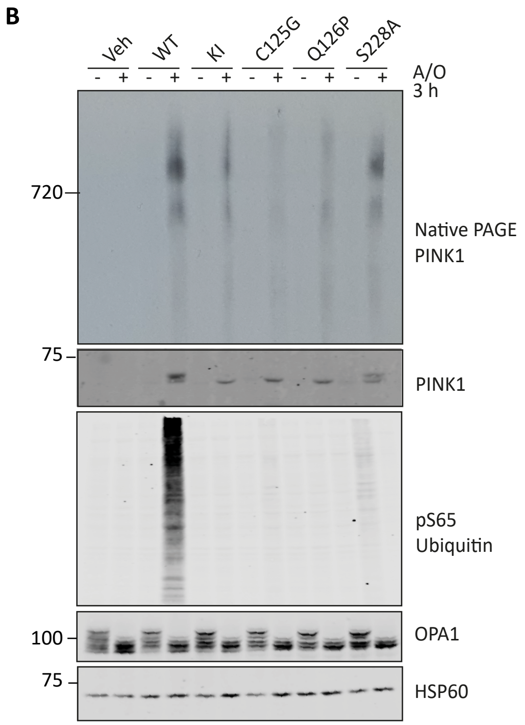
