## Supplementary Table 1 for "Mapping of a N-terminal α-helix domain required for human PINK1 stabilisation, Serine228 autophosphorylation and activation in cells"

| DU Number | **Expresses** | **Type** |
|---|---|---|
| **Control cell lines** | | |
| DU43407 | PINK1 3FLAG (WT) | Full length WT |
| DU46669 | PINK1 3FLAG D384A (KI) | Full length KI |
| DU46848 | PINK1 (111-581)- 3xFLAG | No MTS |
| **Truncation 111-130** | | |
| DU66367 | HSP60 (1-27)-PINK1 104-581-3xFLAG | 104-End |
| DU51362 | HSP60 (1-27)-PINK1 (111-581)-3xFLAG | 111-End |
| DU51855 | HSP60 (1-27)-PINK1 (115-581)-3xFLAG | 115-End |
| DU51817 | HSP60 1-27 PINK1 (120-581) 3xFLAG | 120-End |
| DU51864 | HSP60 (1-27) PINK1 (125-581) 3xFLAG | 125-End |
| DU51812 | HSP60 1-27 PINK1(130-581) 3xFLAG | 130-End |
| **Truncation 111-115** | | |
| DU66370 | HSP60 (1-27)-PINK1 (112-581)-3xFLAG | 112-End |
| DU66368 | HSP60 (1-27)-PINK1 (113-581)-3xFLAG | 113-End |
| DU66382 | HSP60 (1-27)-PINK1 (114-581)-3xFLAG | 114-End |
| **A mutation** | | |
| DU66366 | HSP60 (1-27)-PINK1 111-end-3xFLAG [I111A] | I-A |
| DU66363 | HSP60 (1-27)-PINK1 111-end-3xFLAG [E112A] | E-A |
| DU66381 | HSP60 (1-27)-PINK1 111-end-3xFLAG [E113A] | E-A |
| DU66359 | HSP60 (1-27)-PINK1 111-end-3xFLAG [K114A] | K-A |
| DU66360 | HSP60 (1-27)-PINK1 111-end-3xFLAG [Q115A] | Q-A |
| DU66369 | HSP60 (1-27)-PINK1 111-end-3xFLAG [E117A] | E-A |
| DU66361 | HSP60 (1-27)-PINK1 111-end-3xFLAG [E112A/E113A] | 2E-A |
| DU66373 | HSP60 (1-27)-PINK1-111-581-3xFLAG [E112A/E113A/E117A] | 3E-A |
| **E-K mutations** | | |
| DU66362 | HSP60 (1-27)-PINK1 111-end-3xFLAG [E112K] | E-K |
| DU66365 | HSP60 (1-27)-PINK1 111-end-3xFLAG [E113K) | E-K |

| | | |
|---|---|---|
| DU66528 | HSP60 (1-27)-PINK1 111-end-3xFLAG [E117K] | E-K |
| DU66364 | HSP60 (1-27)-PINK1 111-end-3xFLAG [E112K/E113K] | 2E-K |
| DU66522 | HSP60 (1-27)-PINK1 111-end-3xFLAG [E112K/E113K/E117K] | 3E-K |
| **PD mutations** | | |
| DU66374 | HSP60-(1-27)-PINK1-111-581-3xFLAG (I111S) | I-S |
| DU66521 | HSP60-(1-27)-PINK1-111-581-3xFLAG (Q115L) | Q-L |
| DU66376 | HSP60-(1-27)-PINK1-111-581-3xFLAG (Q126P) | Q-P |
| DU66375 | HSP60-(1-27)-PINK1-111-581-3xFLAG (C125G) | C-G |
| DU56048 | pcDNA5 FRT/TO Pink1 3FLAG C125G | C-G |
| DU56186 | pcDNA5 FRT/TO Pink1 3FLAG Q126P | Q-P |
| DU56077 | pcDNA5 FRT/TO Pink1 3FLAG A168P | A-P |
| DU56079 | pcDNA5 FRT/TO Pink1 3FLAG E240K | E-K |
| DU56049 | pcDNA5 FRT/TO Pink1 3FLAG G309D | G-D |
| DU56085 | pcDNA5 FRT/TO Pink1 3FLAG G409V | G-V |
| DU56537 | pcDNA5 FRT/TO Pink1 3FLAG L539F | L-F |
| DU67934 | pcDNA5 FRT TO PINK1 534_535InsQ 3FLAG | 534_535insQ |
| **TOM 20 tethering MTSs** | | |
| DU66429 | pOTC-(1-33)-PINK1-(111-581)-3xFLAG | WT MTS |
| DU66430 | pOTC-(1-33)-PINK1-(111-581)-3xFLAG (L5A, L8A, L9A) | Unable to bind TOM20 |
| DU66451 | F1 Beta-ATPase-(1-33)-PINK1-(111-581)-3xFLAG | WT MTS |
| DU66431 | F1 Beta-ATPase-(1-33)-PINK1-(111-581)-3xFLAG (W29A, C32A, M33A) | Unable to bind TOM20 |
| DU66540 | PINK1-M1-P34, I111-END-3xFLAG | 35-110 deletion |
| Disease mutants | | |
| **CTE mutants** | | |
| DU27429 | PINK1-3FLAG pcDNA5 FRT/TO (V528A) | V-A |
| DU60932 | PINK1-3FLAG pcDNA5 FRT/TO (L532A) | L-A |
| DU60929 | PINK1-3FLAG pcDNA5 FRT/TO (L539A) | L-A |

| | | |
|---|---|---|
| DU67239 | PINK1-3FLAG pcDNA5 FRT/TO (V528A / L532A / L539A) | 3A |
| **Pediculus mutations** | | |
| DU66324 | PhPINK1 (108-end) | |
| DU66537 | PhPINK1 (108-end) L507A | L-A |
| DU66538 | PhPINK1 (108-end) L514F | L-F |
| DU66536 | PhPINK1 (108-end) I503A | I-A |
| DU72026 | PhPINK1 (108-end) W129P | W-P |
| DU66535 | PhPINK1 (108-end) C128G | C-G |
| DU72025 | PhPINK1 (108-end) E125K | E-K |
